# A SRC-Annexin A2 axis that couples membrane repair to microRNA export during radiation stress in glioblastoma

**DOI:** 10.64898/2026.04.21.719868

**Authors:** Shilpi Singh, Akhilesh Kumar Maurya, Iteeshree Mohapatra, Stefan Kim, Afsar R. Naqvi, Gatikrushna Singh, Clark C. Chen

## Abstract

Repair of ionizing-radiation (IR)-induced membrane damage is essential for cell survival, and Annexin A2 (ANXA2) mediates this repair by promoting shedding of microvesicles containing both damaged lipid and ANXA2. Here, we show that coupling this process to microRNA export contributes to radiation resistance of glioblastoma, the most common adult brain cancer. Structural modeling and mutational analysis identify that ANXA2 tyrosine 23 (Y23) phosphorylation, required for its IR-induced localization to damaged membranes, is also essential for binding miR-603, a critical regulator of glioblastoma cell state. This binding is required for IR-induced miR-603 export, indicating that the repair program recruiting ANXA2 also positions it to load miR-603 into microvesicles. Mutations that abolish ROS-induced sulfenylation and activation of SRC (SRC-C185A/C245A) suppressed ANXA2-Y23 phosphorylation, microvesicle shedding, and miR-603 export. The epistatic interaction between ANXA2-Y23A, SRC-C185A/C245A, and miR-603 delineates a therapeutically targetable radiation-response network that couples membrane repair with miRNA export.

## INTRODUCTION

Glioblastoma is the most common and lethal primary brain tumor in adults, marked by rapid progression and near-universal recurrence^1^. Ionizing radiation (IR) remains one of the few treatments with proven survival benefit, yet its efficacy is ultimately limited by the near-inevitable development of acquired resistance^2^. This resistance reflects a multifaceted adaptive program involving a wide range of cellular processes that converge on an integrated stress-response network that enables cellular adaptation to diverse stress conditions^3^. Within this network, microRNA homeostasis serves as a key regulatory layer^4^; these small, non-coding RNAs fine-tune gene-expression programs that govern cellular stress-adaptation^5^.

Stress conditions, including exposure to ionizing radiation (IR), alter miRNA homeostasis to promote adaptation^6^. A growing body of evidence indicates that stress-induced miRNA export is a major mechanism for modulating miRNA homeostasis during stress adaptation^7,8^. In glioblastoma, the most common form of primary brain cancer in adults^1^, IR treatment induces cellular export of miR-603 through the release of extracellular vesicles (EVs)^9^. Cellular depletion of miR-603, in turn, de-represses its downstream targets, including insulin-like growth factor 1 (IGF1) and its cognate receptor (IGF1R)^9,10^. The subsequent surge in IGF signaling promotes the transition to stem-like cell states that drive radiation resistance^9^. The specific EV subtypes that mediate this export process, as well as the underlying molecular machinery, have not yet been defined. This lack of mechanistic clarity represents a significant gap in our understanding of acquired radiation resistance.

EVs are a heterogeneous population of membrane-bound particles released by cells that contribute to cellular homeostasis and mediate intercellular communication^11,12^. They represent a major class of extracellular macromolecular particulates, comprising a diverse array of nano- to micro-scale particles released into the extracellular space^8,9^. EVs are distinguished from other macro-particulates by their membrane-enclosed vesicular structure and the characteristic surface markers displayed on that membrane. EVs lack apolipoprotein A1 (ApoA1) and Annexin V, which characterize lipoproteins and apoptotic bodies, respectively^13^. EVs are further subclassified based on vesicle size, surface protein expression, and cellular origin^14,15^. Exosomes, defined as EVs <150 nm, are enriched in tetraspanins such as CD63, CD9, and CD81 on their membranes^16^. EVs >150 nm are termed microvesicles (MVs) and are characterized by enrichment of surface proteins, including integrin alpha-V^17^, calnexin^18,19^, and HSPA5^20^.

IR exposure induces a robust increase in EV biogenesis and secretion^21^. The burst of reactive oxygen species (ROS) generated by the ionization of water molecules^22^ causes lipid peroxidation, disrupting the structural integrity of the phospholipid bilayer^23^. Radiation-induced damage to the plasma membrane causes uncontrolled ion flux, including calcium (Ca²⁺) influx^24,25^. This rise in intracellular Ca²⁺ recruits Annexin A2 (ANXA2), a calcium-dependent phospholipid-binding protein, to sites of exposed acidic lipids^26,27^. There, it forms a heterotetramer with S100A10^28,29^, generating a scaffold that stabilizes the damaged plasma membrane. The Ca²⁺ influx subsequently activates calpain, a calcium-activated cysteine protease^30,31^, which proteolyzes ANXA2 at defined sites to facilitate outward budding and excision of the damaged membrane^32^, with ANXA2 incorporated into the resulting microvesicle. Of note, higher ANXA2 expression in glioblastoma consistently correlated with poor prognosis across independent cohorts^33–36^.

Cell-surface translocation of ANXA2 requires phosphorylation of its tyrosine 23 (Y23) residue^36–38^, a modification governed by steroid receptor coactivator (SRC) family serine and tyrosine kinases^37,38^. SRC itself is a redox-sensitive kinase that functions as a sensor of oxidative stress via reversible cysteine sulfenylation^39,40^. Hydrogen peroxide (H₂O₂)-induced ROS modify cystine (Cys185 and Cys277) residues of SRC, generating sulfenylated intermediates (Cys-SOH). These sylfenylation events destabilize the autoinhibited conformation of SRC, shifting it toward an active state^41^. Activated SRC, in turn, stimulates NADPH oxidase^42,43^, generating additional ROS and creating a feed-forward redox loop^44^. Comparative structural analysis revealed that multiple SRC-family kinases share homologous redox-active cysteines, suggesting that oxidative activation is a generalizable regulatory mechanism^40,41^.

In this study, we investigated the mechanism underlying IR-induced export of miR-603 and identified MVs as the responsible mediator. Proteomic profiling of biotinylated miR-603 pull-down from these MVs identified ANXA2 as a miR-603-binding partner. This binding requires Y23 phosphorylation of ANXA2 and is abolished in SRC C185A and C277A mutants. Perturbations impairing ANXA2 expression or ANXA2 Y23 phosphorylation suppressed IR-induced MV release and export of miR-603-containing MVs, thereby enhancing radiation sensitivity. These findings support a model in which SRC and ANXA2 coordinate radiation-induced membrane repair and miRNA export, contributing to acquired radiation resistance.

## RESULTS

### Ionizing radiation (IR) induces microvesicle (MV)-mediated export of miR-603

To identify the EV population responsible for IR-induced miR-603 export, we transfected Cy5-labeled miR-603 into LN340 stably expressing CD63-GFP^16^, allowing visualization of GFP-labeled exosomes. This approach enabled us to determine whether Cy5-miR-603 is released within CD63-positive exosomes in response to IR treatment. While IR induced robust Cy5-miR-603 export via EV release, these Cy5-miR-603-containing vesicles were CD63-GFP–negative and notably larger than the expected size of exosomes (**Fig. 1A**, right lower panel; video shown in **Fig. S1**). Quantitative analysis across 100 imaging fields showed that IR induced a ∼10-fold increase in the release of Cy5-miR-603 containing, large vesicles (**Fig 1A**, right panel bar graph).

**Figure 1:**
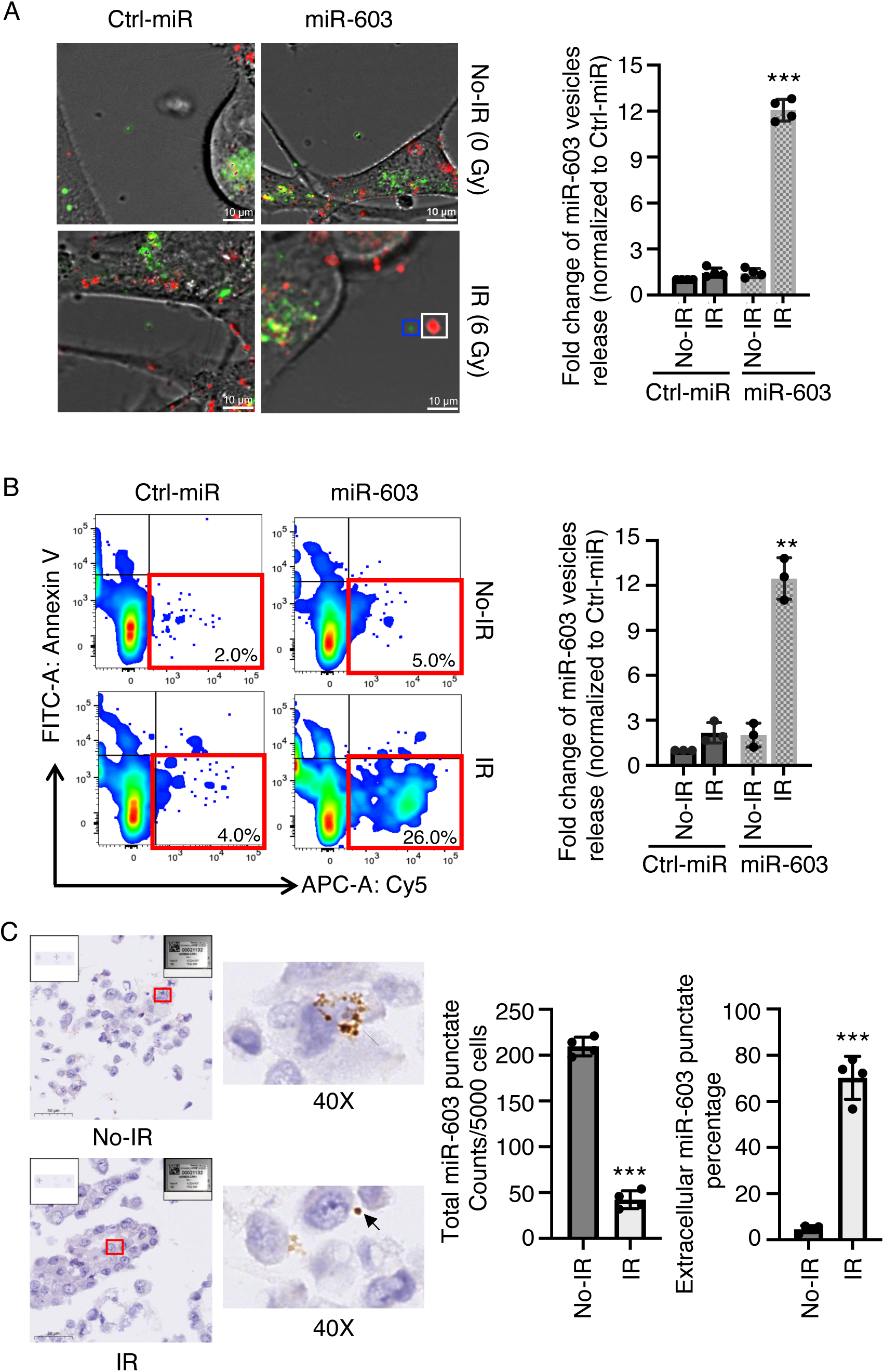
Ionizing radiation (IR) induces microvesicle (MV)-mediated export of miR-603. **A.** Cy5-miR-603 is released in CD63⁻ extracellular vesicles (EVs) larger than exosome. CD63-GFP expressing LN340 was transfected with Cy5-tagged miR-603 or miR-NT (Ctrl-miR) and treated with 6 Gy IR or mock treatment and monitored using live-cell fluorescence imaging. *<u>Left</u>*: representative images of EV release (actual video shown in **Fig. S1A**). Blue box: CD63-GFP^+^ EVs; white box: Cy5-miR-603^+^ EVs. *<u>Right</u>*: Quantification of Cy5-miR-603^+^ EVs across 100 fields. ***p < 0.001 between indicated groups (Student’s t-test). Scale bar is 10 μm. **B.** miR-603 is exported in Annexin V^-^ vesicles. EVs are isolated from LN340 cells transfected with Cy5-tagged miR-603 or miR-NT (Ctrl-miR), treated with 6 Gy IR or mock treatment, and analyzed by flow cytometry. *<u>Left</u>*: Cy5/FITC-Annexin V contour plot. Red box: Cy5^+^/Annexin V^-^ EVs. *<u>Right</u>*: Quantification Cy5^+^/Annexin V^-^ EVs. ***p < 0.001 between indicated groups (Student’s t-test). **C.** Radiation induced export of endogenous miR-603. LN340 cells were treated 6 Gy IR or mock treatment, cultured for 12, and processed for RNAscope HD assay for miR-603. *<u>Left</u>*: representative images of nuclei counterstained with hematoxylin. Brown punctate signals indicate miR-603 detection. *<u>Right</u>*: the number of cytoplasmic and extracellular miR-603 puncta quantified from 5,000 cells treated with or without IR (left). The percentage of extracellular miR-603 puncta is shown (right). Data represent mean ± SD. ***p< 0.001.

To determine whether the large Cy5-miR-603-containing vesicles were apoptotic bodies^44^, we isolated the total extracellular contents from LN340 conditioned media after IR or mock treatment by centrifugation and assessed for annexin V staining using flow cytometry (**Fig. 1B**). IR increased the abundance of annexin V-positive (+) particles, consistent with apoptotic body formation, but most of the Cy5-miR-603 vesicles were annexin V-negative (-), indicating most miR-603 containing EVs are unlikely apoptotic bodies (**Fig. 1B**, right lower panel indicated in red box and bar graph).

To exclude artifacts related to Cy5-miR-603 transfection, we next assessed endogenous miR-603 using RNAscope (Advanced Cell Diagnostics) in situ hybridization^45,46^. This approach enabled visualization of native miR-603 without transfection of exogenous miRNAs. In non-irradiated LN340 cells, miR-603 was predominantly confined to intracellular compartments, appearing as discrete cytoplasmic puncta (**Fig. 1C**, upper panels). IR treatment induced a redistribution of miR-603, with punctate accumulation in the extracellular space (**Fig, 1C**, lower panels). Quantitative analysis across 100 fields showed that IR reduced intracellular miR-603 puncta by approximately fourfold and increased extracellular miR-603 by approximately two orders of magnitude (**Fig. 1C**, bar graphs). Together, these findings support the hypothesis that IR induces miR-603 export via a non-exosomal, non-apoptotic EV population.

### Characterization of miR-603-containing MVs

Next, we characterized the large vesicles containing miR-603 that are released in response to IR. LN340 cells were transfected with Cy5-labeled miR-603 and exposed to IR. The conditioned media was collected, and cell debris were removed by low-speed centrifugation followed by total EVs isolation at > 100,000 g centrifugation (**Fig. 2A**). Nanoparticle tracking analysis (NTA) of the EVs showed IR-induced an increase in vesicles >150 nm in size (hereto referred as MV fraction, **Fig. 2A**, red arrow), consistent with literature demonstrating that IR promotes MV shedding^47,48^. The abundance of vesicles <150 nm was not significantly altered by IR treatment (**Fig. 2A**). The total EV pool was subsequently fractionated by differential ultracentrifugation to isolate EVs that were enriched for MVs (10,000 g pellet) or exosomes (100,000 g pellet) (**Fig. S2**). The pellets of these centrifugation runs were washed, stained with Annexin V, and analyzed by flow cytometry for Cy5-containing EVs. In this analysis, IR treatment increased Cy5-containing EVs only within the MV fraction (**Fig. 2B**, represented in red **and S3,** upper panels). In contrast, the exosome fractions were largely Cy5 negative (**Fig. 2B**, represented in blue). Notably, flow-cytometric analysis of vesicles within the MV fraction indicates that they are mostly Cy5-positive (+) and Annexin V-negative (-) (**Fig. S3**).

**Figure 2.**
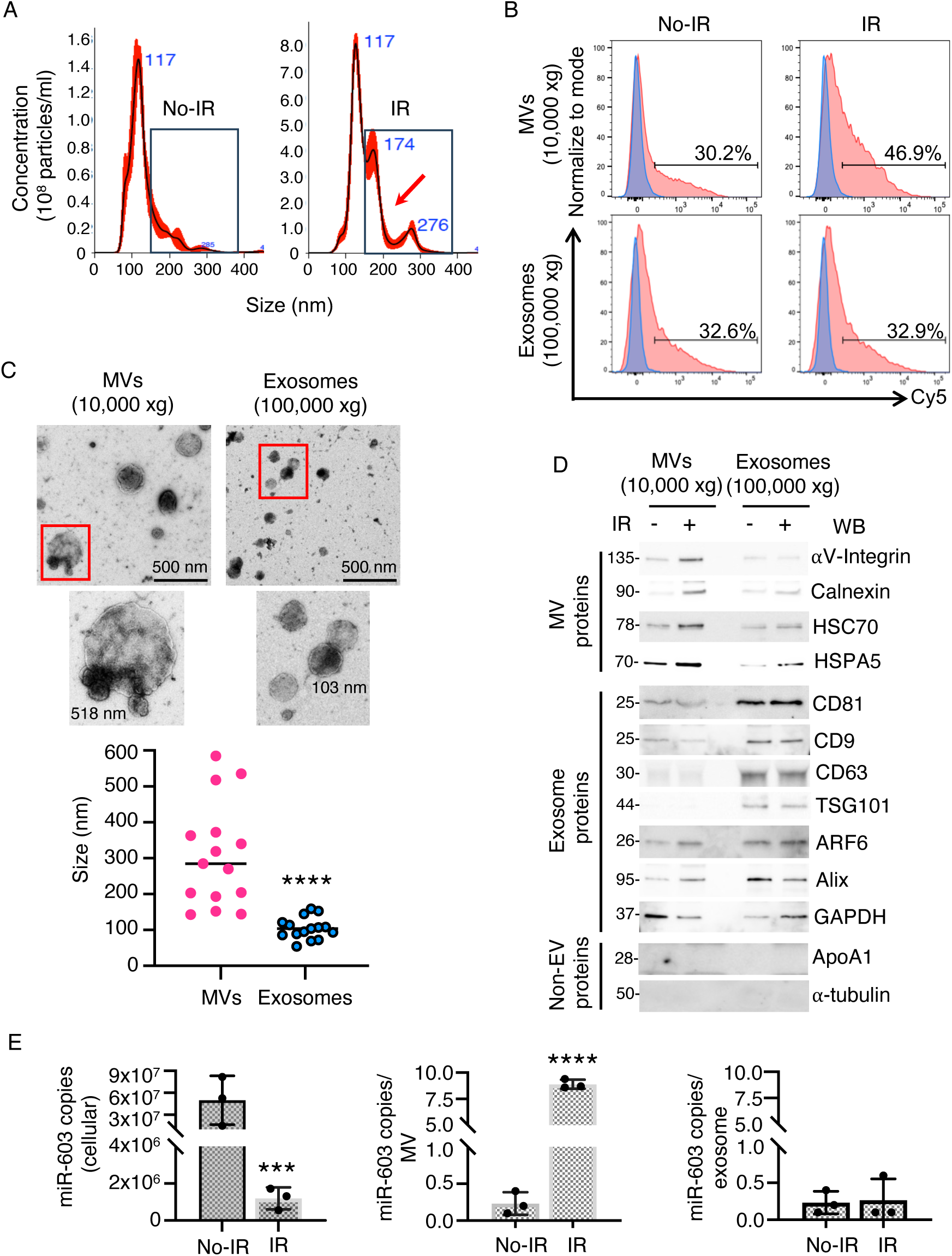
Characterization of miR-603-containing microvesicles (MVs). **A.** Radiation induced an increase in the release of vesicles > 150 nm. Nanosight nanoparticle tracking analysis of conditioned media isolated from LN340 cells treated with 6 Gy IR or mock treatment. Representative nanoparticle tracking shows the distribution of particle size as a function of particle number (particles/mL) and modal size (nm). The box indicates particles > 150 nm (enriched for microvesicles (MVs)). **B.** Differential ultracentrifugation–based isolation of MVs (10,000 × g) and exosomes (100,000 × g) from media depleted of cellular debris. MVs and exosomes were isolated from Cy5-miR-603 or Cy5-miR-Ctrl transfected LN340 cells (with or without radiation treatment) and subjected to Cy5/FITC-Annexin V flow cytometry. Each treatment group is normalized to the mode and plotted in histogram (peak=100%). Blue: vesicles from the cells with Cy5-miR-Ctrl-transfection. Red: vesicles from the cells transfected with Cy5-miR-603. **C.** Transmission electron micrograph of <150 nm (exosome-enriched) and >150 nm (MV-enriched) EV’s isolated from irradiated LN340 by differential centrifugation and negative staining of the vesicles. Scale Bar: 500 nm. Image magnification 18.5 k. Lower panel: EVs size distribution by TEM analysis. Mann-Whitney test: ****p < 0.0001. **D.** Immunoblot analysis of EV markers on <150 nm (exosome-enriched) and >150 nm (MV-enriched) EV’s isolated from non-irradiated and irradiated LN340 cells condition media. **E.** RT-qPCR analysis of miR-603 levels in cytoplasmic, MV, and exosome fractions. LN340 cells were treated with 6 Gy IR or mock control. RNA was extracted from cells as well as from MVs and exosomes isolated from conditioned media, and miR-603 levels were quantified by RT-qPCR.

To confirm the size distribution of the two EV subsets, vesicles were analyzed by transmission electron microscopy (TEM) upon negative staining (**Fig. 2C**). In this analysis, the sizes of the EVs in the MV fraction ranged from ∼200-600 nm in diameter (**Fig. 2C**, left and bottom panel), whereas the EVs in the exosomes ranged from ∼50-150 nm (**Fig. 2C**, right and bottom panel).

Next, we characterized the protein contents of EVs in the MV and exosome fractions by Western blotting. The EVs in the MV (10,000 g) fraction showed an enrichment of integrin-αV, calnexin, and HSPA5, and the absence of the canonical exosomal markers CD63, CD9, and CD81 (**Fig. 2D**). The levels of the MV-associated proteins increased in the MVs isolated after IR. In contrast, EVs in the exosome (100,000 g) fraction showed an enrichment of CD63, CD9, and CD81. ApoA1 was not detected in either EV fraction. The levels of exosome-associated proteins in the exoome fraction remain unchanged after IR treatment. Together, these findings suggest that IR drives the release of miR-603 through large vesicles bearing a characteristic MV surface marker profile and deficient in ApoA1.

Next, MVs and exosomes were isolated from conditioned media of irradiated LN340 cells, and the miR-603 copy number was quantified in MV-enriched fractions, exosome-enriched fractions, and matched cellular lysates. In this experiment, IR induced a five-fold reduction in cellular miR-603 (**Fig. 2E**, left panel), accompanied by an approximately ten-fold increase in MV-associated miR-603 (**Fig. 2E**, middle panel). No notable changes in miR-603 levels were observed in the exosome fraction (**Fig. 2E**, right panel). These findings were consistently reproduced across independent glioblastoma cell lines (**Fig. S4**). Together, these findings suggest that IR induces the export of miR-603 through MV release.

### ANXA2 is necessary for IR- induced MV mediated miR-603 export

To define the mechanism underlying IR-induced miR-603 export, we sought to identify the MV protein that binds miR-603. To this end, MVs were isolated from irradiated and non-irradiated LN340 cells transfected with biotinylated-miR-603 (Bi-miR-603) or a biotinylated-control-miRNA (Bi-miR-ctrl). The biotinylated RNAs were affinity-purified, and associated proteins were subsequently eluted and subjected to silver staining (**Fig. S5A**) and mass-spectrometry–based proteomic profiling (see **Methods**). Of note, a ∼39 kDa band is uniquely present in the Bi-miR-603–transfected, IR-treated sample (**Fig. S5A**). Top candidates in each of the four conditions (Bi-miR-ctrl pull-down from non-irradiated and irradiated cells, Bi-miR-603 pull-down from non-irradiated and irradiated cells) were cross-referenced to identify those found only in the Bi-miR-603 pull-down from irradiated cells (boxed in red in **Fig. S5B,** light blue: indicates absence of a protein, and pink indicates presence of a protein in the proteomic profile).

Among the candidates, annexin 2 (ANXA2) stood out as a ∼39 kDa protein central in the repair of IR-damaged membranes. ANXA2 is recruited to IR-damaged membranes, where it stabilizes injured membrane domains, facilitates the budding of damaged fragments through MV release, and is itself shed in these MVs^49^. These findings raise the possibility that ANXA2 chaperones miR-603 to the plasma membrane during repair and facilitates miRNA export. To confirm miR-603 ANXA2 interaction, we transfected Bi-miR-603 or Bi-miR-ctrl into LN340 cells. The transfected cells were treated with IR or mock treatment. MVs were isolated, and the lysates were subjected to streptavidin affinity pulldown followed by ANXA2 immunoblotting. The result showed a notable ANXA2 enrichment in the Bi-miR-603 pulldown from irradiated cells, relative to all other conditions (**Fig. 3A**). Reciprocal immunoprecipitation using an anti-ANXA2 antibody was performed on the MVs lysate, followed by RNA extraction from the immunoprecipitates, and miR-603 RT-qPCR. This experiment revealed an order-of-magnitude enrichment of miR-603 copies in the immunoprecipitates from irradiated cells (**Fig. 3B**). Together, these findings suggest that ANXA2 associates with miR-603 in an IR-dependent manner.

**Figure 3.**
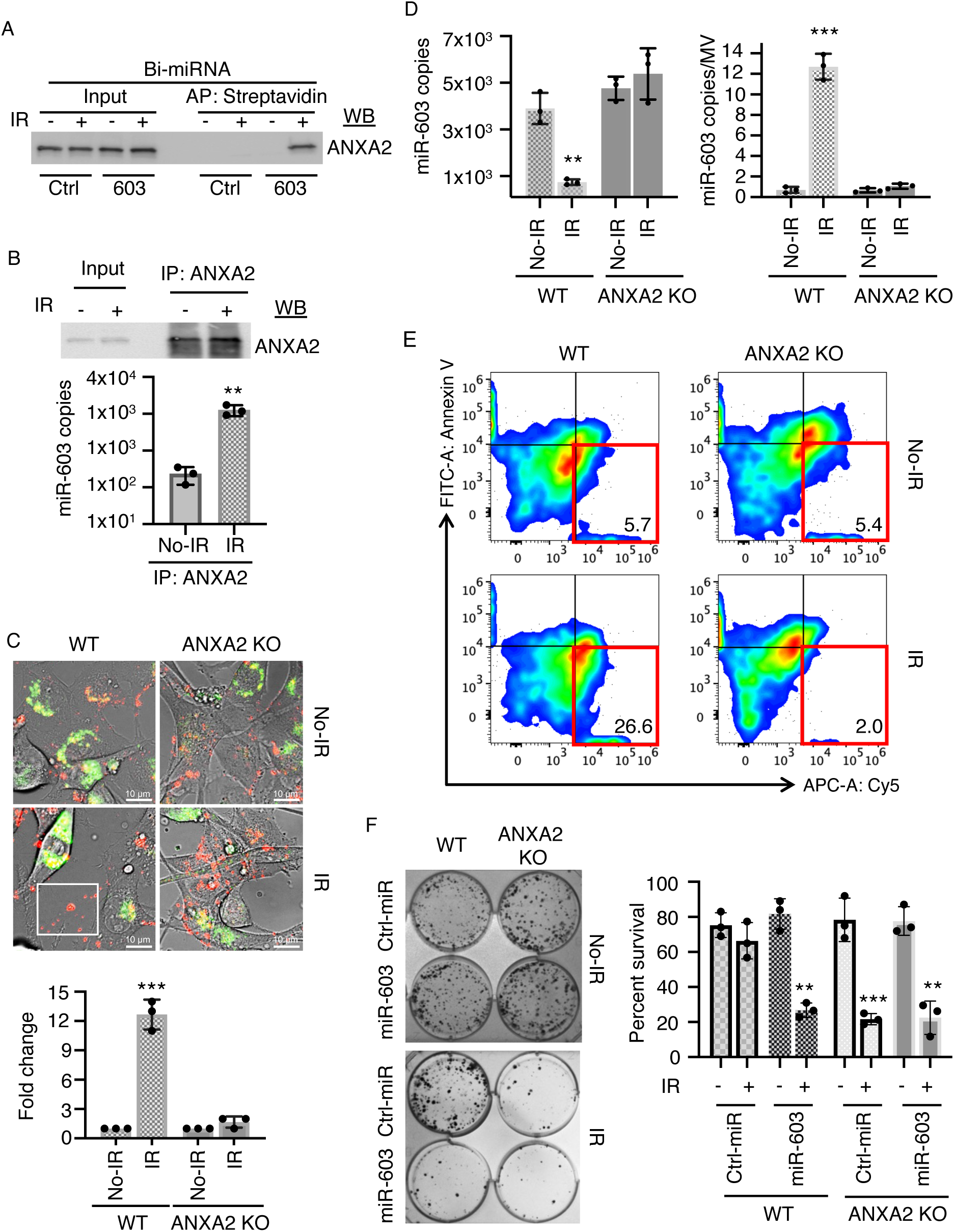
Interaction between miR-603 and Annexin A2 (ANXA2). **A.** ANXA2 interacts with biotinylated-miR-603 (Bi-miR-603) in a radiation-dependent manner. MVs were isolated from irradiated and non-irradiated LN340 cells transfected with Bi-miR-603 or a Bi-miR-Ctrl. The biotinylated RNAs were subjected to streptavidin pulldown followed by immunoblotting for ANXA2. **B.** miR-603 is enriched in the ANXA2 immunoprecipitation (IP). MVs were isolated from above and subjected to ANXA2 IP, followed by RNA isolation from the IPed fractions and RT-qPCR for miR-603. *<u>Top</u>*: ANXA2 immunoblotting of the IPed fractions. *<u>Bottom</u>*: miR-603 RT-qPCR of RNA isolated from the same fractions. **C.** ANXA2 is required for radiation-induced export of miR-603-containing MVs. Wild-type (WT) and CRISPR ANXA2-knockout (KO) LN340 CD63-GFP cells were transfected with Cy5-miR-603, subjected to irradiation or mock treatment, and monitored by live-cell imaging. *<u>Top</u>*: representative images of vesicle release (actual video shown in **Fig. S6B**). White box: Cy5-miR-603^+^ EVs. *<u>Bottom</u>*: Quantification of Cy5-miR-603^+^ EVs across 100 fields. ***p < 0.001 between indicated groups (Student’s t-test). Scale bar is 10 μm. **D.** ANXA2 is required for the IR-induced shift of miR-603 from the cytoplasm to MVs. Total RNA was isolated from WT and ANXA2 KO LN340 cells following 6 Gy IR or mock treatment, followed by miR-603 RT-qPCR. Data represent mean ± SD. **p< 0.01 and ***p< 0.001. **E.** ANXA2 required for miR-603 is exported in Annexin V^-^ vesicles. EVs are isolated from LN340 cells transfected with Cy5-tagged miR-603, treated with 6 Gy IR or mock treatment, and analyzed by flow cytometry. Cy5/FITC-Annexin V^-^contour plot for the *<u>Left</u>*: MVs isolated from WT cells transfected with Cy5-miR-603. *<u>Right</u>*: MVs isolated from ANXA2 KO cells transfected with Cy5-miR-603. Red box: Cy5^+^/Annexin V^-^ EVs. **F.** Epistatic interaction between ANXA2 and miR-603 in radiation sensitivity modulation. WT and ANXA2 KO LN340 cells were transfected with miR-603 or control miR and subjected to clonogenic survival analysis. *<u>Left</u>*: representative clonogenic experimental image. *<u>Right</u>*: quantitation of clonogenic survival. Bar graphs represent the mean ± SD of percent survival. **p< 0.01.

Next, we determined whether ANXA2 is necessary for IR-induced miR-603 export. To this end, ANXA2 was disrupted using CRISPR-mediated gene knockout. Consistent with published literature^50^, the ANXA2-knockout (KO) showed increased IR sensitivity and accumulation of γ−H2AX in response to IR (**Fig. S6A**). Wild-type (WT) and CRISPR ANXA2-knockout (KO) LN340-CD63-GFP cells were then transfected with Cy5-miR-603, subjected to irradiation or mock treatment, and monitored by live-cell imaging (**Fig. 3C**, video in **Fig. S6B**). IR induced more than a 10-fold increase in Cy5-miR-603-containing MVs release in the WT LN340 but not in ANXA2 KO LN340 cells (**Fig. 3C**, bar graph). Further supporting the requirement for ANXA2 in miR-603 export, IR induced an ∼3-fold reduction in cytoplasmic miR-603 (**Fig. 3D**, left panel) and an ∼10-fold increase in MV-associated miR-603 in WT cells (**Fig. 3D**, right panel). Both changes were absent in the ANXA2-KO cells (**Fig. 3D**). To further confirm the role of ANXA2 in miR-603 export, MVs were isolated from the conditioned media of Cy5-miR-603-transfected LN340 WT or ANXA2 KO cells after IR or mock treatment. Flow cytometric analysis of these MVs showed that the proportion of Cy5⁺/Annexin V⁻ MVs was reduced by an order of magnitude in ANXA2 KO cells relative to WT LN340 (**Fig. 3E**). Collectively, the data support ANXA2 as an essential mediator of the IR-driven MV release and miR-603 export.

ANXA2 forms a stable heterotetrameric complex with S100A10 that is essential for membrane trafficking and Ca^2+^-regulated membrane repair^51,52^. To evaluate whether S100A10 contributes to radiation-induced miR-603 export, LN340 cells were transfected with S100A10-targeting siRNA and exposed to IR. MVs were isolated from conditioned media, and miR-603 levels were quantified by RT-qPCR. In cells transfected with control siRNA (si-NT), IR induced a ∼5-fold decrease in cytoplasmic miR-603, accompanied by a ∼5-fold increase in MV-associated miR-603. These changes were abolished upon S100A10 silencing (**Fig. S7**), mirroring the phenotype observed in ANXA2 knockout cells (**Fig. 3D**).

Previous studies indicate that loss of ANXA2 increases glioblastoma radiosensitivity^53^. Our findings support a model in which ANXA2 exerts this effect through miR-603 export, a hypothesis that predicts the radiation-sensitizing effects of ANXA2 loss and miR-603 restoration would be epistatic. Supporting this hypothesis, ANXA2 KO and miR-603 transfection each produced a three-fold increase in radiation sensitivity of WT LN340 (**Fig. 3F**). miR-603 transfection into the ANXA2 KO cells did not further enhance radiosensitivity, demonstrating epistasis (**Fig. 3F**). Further supporting this epistasis, transfection of miR-603 in WT LN340 increased comet tail accumulation in response to IR (**Fig. S8A and B**), and this induction was absent in ANXA2-KO LN340 cells. These findings suggest that loss of ANXA2 elevates cytoplasmic miR-603 to a level sufficient to suppress DNA repair, such that additional, exogenously introduced miR-603 produces no further effect on this phenotype. Collectively, our findings establish that ANXA2 is necessary for miR-603 export and that its impact on radiation sensitivity is mediated through this regulatory mechanism.

### Tyrosine 23 phosphorylation of ANXA2 is required for miR-603 binding

ANXA2 is known to undergo phosphorylation at multiple residues, the most well-characterized of which include Ser11, Tyr23, and Ser25^54–57^. Of note, phosphorylation of tyrosine 23 (Y23) is essential for ANXA2 recruitment to IR-damaged membranes and for its role in stabilizing and repairing injured membranes^58,59^. We next determined whether mutations that disrupt these phosphorylation sites impaired the interaction between ANXA2 and miR-603. To this end, ANXA2 S11A or Y23A or S25A mutants were expressed in ANXA2-KO LN340 cells, followed by the transfection of Bi-miR-603 and irradiation. MVs were isolated from the conditioned media, and the lysates were subjected to Bi-miR-603 pulldown. ANXA2 immunoblotting of these pulldowns revealed a loss of ANXA2 only in the Bi-miR-603 pulldown from the Y23A mutant (**Fig. 4A**). Reciprocal immunoprecipitation using an α-ANXA2 antibody was performed on the MV lysate from ANXA2 KO LN340 cells expressing ANXA2 S11A, Y23A, or S25A mutants. RNA was extracted from the immunoprecipitated and subjected to miR-603 RT-qPCR. This experiment revealed more than an order-of-magnitude enrichment of miR-603 in ANXA2 immunoprecipitates from irradiated cells transfected with WT ANXA2, S11A, or S25A. This enrichment was not observed in immunoprecipitates derived from ANXA2 KO cells transfected with Y23A mutants. These results suggest that Y23 phosphorylation is critical for the interaction between miR-603 and ANXA2 (**Fig. 4B**).

**Figure 4.**
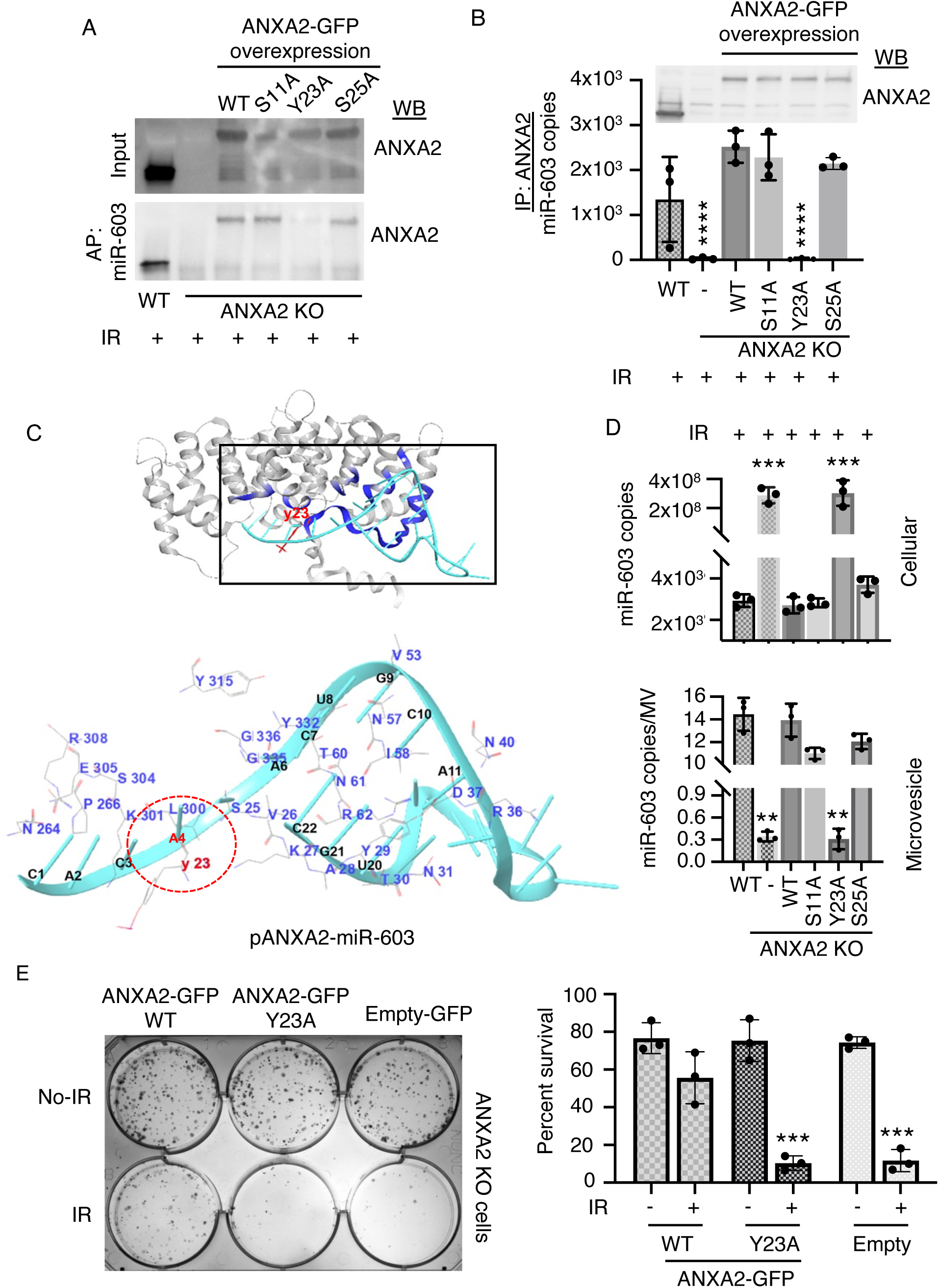
Tyrosine 23 phosphorylation of ANXA2 is required for miR-603 binding. **A.** Mutation of ANXA2 Tyr23 (Y23) abolishes binding to biotinylated miR-603. ANXA2-KO LN340 cells were transfected with ANXA2 mutants (S11A, Y23A, or S25A) for 24 h, followed by Bi-miR-603 transfection and 6 Gy irradiation. MVs isolated from conditioned media were lysed and subjected to streptavidin pulldown of Bi-miR-603, and bound proteins were detected by ANXA2 immunoblotting. **B.** miR-603 is not enriched in ANXA2 immunoprecipitates from MVs of ANXA2-Y23A-expressing cells. MVs were isolated as described in (A) and subjected to ANXA2 immunoprecipitation, followed by RNA extraction and RT-qPCR analysis of miR-603. *<u>Top</u>*: ANXA2 immunoblot of immunoprecipitated samples. *<u>Bottom</u>*: miR-603 RT-qPCR of RNA isolated from the immunoprecipitates. Data represent mean ± SD. ****p< 0.0001. **C.** Molecular Dynamics (MD) simulation identified high-confidence interaction models for miR-603 with pANXA2. pANXA2 structure shown in gray and blue. miR-603: cyan ribbon; miR-603 interacting region of ANXA2 shown in blue. ANXA2 residues: red/gray stick. The red circle indicates Y23 residue of ANXA2. miR-603 nucleotides shown in black. The red dotted circle indicates the interaction of phosphorylated Y23 of ANXA2 and A4 of miR-603. **D.** ANXA2 Y23A mutation abolishes IR-induced packaging of cytoplasmic miR-603 into MVs. Total RNA was isolated from WT and ANXA2 KO LN340 cells and their corresponding MVs after 6 Gy IR treatment, followed by RT-qPCR quantification of miR-603. Data represent mean ± SD. ***p < 0.001. **E.** Y23A mutant did not rescue the radiation resistance in ANXA2 KO cells. LN340 and ANXA2 KO cells were transfected with WT ANXA2 or the Y23A mutant plasmids followed by 0 or 6 Gy IR and subjected to clonogenic survival analysis. *<u>Left</u>*: representative clonogenic experimental image. *<u>Right</u>*: quantitation of clonogenic survival. Bar graphs represent the mean ± SD of percent survival. ***p< 0.001.

The availability of the ANXA2 crystal structure enabled molecular dynamics (MD) simulations to examine how Y23 phosphorylation influences miR-603 binding (**Fig. 4C and S9**). Docking analysis identified a high-confidence interaction model between miR-603 and phosphorylated ANXA2 (pANXA2). Phosphorylated Y23 interacts with the adenine residue at position 4 (A4) of miR-603 (**Fig. 4C**). This critical interaction is absent in unphosphorylated ANXA2 (**Fig. S9A**), and the alanine substitution mutation at ANXA2-Y23A abolished this interaction (**Fig. S9B**). Molecular dynamics (MD) analysis further revealed that the ANXA2-miR-603 complex stabilized rapidly (∼20 ns), consistent with a relatively constrained structure. pANXA2 exhibited increased conformational flexibility, with sustained RMSD fluctuations up to ∼50 ns. Binding of miR-603 to pANXA2 stabilized this dynamic state, as evidenced by the pANXA2-miR-603 complex exhibiting minimal RMSD drift (**Fig. S10A**). Energetic analysis further revealed that the pY23 ANXA2-miR-603 complex exhibits stronger Coulombic electrostatic interactions with minimal temporal fluctuations relative to the ANXA2-miR-603 complex (**Fig. S10B**).

Free-energy landscape (FEL) analysis provided additional insight into conformational stability. While ANXA2 alone sampled multiple shallow energy basins (**Fig. S10C**) and the ANXA2-miR-603 complex occupied two low-energy states (**Fig. S10D**), the pANXA2-miR-603 complex converged into a single deep energy basin, indicating strong conformational stabilization upon RNA binding (**Fig. S10E and F**). In contrast, the ANXA2-Y23A-miR-603 complex failed to form a stable low-energy state, consistent with dynamic instability (**Fig. S10G and H**). Collectively, these analyses demonstrate that phosphorylation of ANXA2 at Y23 enhances electrostatic and van der Waals interactions with miR-603, stabilizes the protein-RNA complex, and promotes a favorable binding energy landscape. Loss of this modification due to the Y23A mutation disrupts these interactions, leading to reduced binding affinity and structural instability. These findings support a model in which Y23 phosphorylation of ANXA2 is required for efficient miR-603 binding.

We next determined if Y23 phosphorylation is required for IR-induced MVs release and miR-603 export. Nanoparticle tracking analysis of EVs isolated from conditioned media of ANXA2-KO cells transfected with ANXA2 Y23A and exposed to IR showed a significant reduction in the number of particles > 150 nm (**Fig. S11**, lower middle panel). Such a reduction was not seen in cells transfected with WT, S11A, or S25A ANXA2 plasmid (**Fig. S11**, left and right panels). Further supporting the requirement for Y23 phosphorylation in miR-603 export, ANXA2 KO cells transfected with WT, S11A, or S25A ANXA2 exhibited the IR-induced reduction in cytoplasmic miR-603 (**Fig. 4D**, top panel) and corresponding increase in MV-associated miR-603 (**Fig. 4D**, bottom panel); these changes were absent in cells transfected with the Y23A mutant (**Fig. 4D**). Finally, the Y23A mutant failed to rescue the radiation sensitivity of ANXA2 KO cells (**Fig. 4E**, middle well and bar graph), whereas transfection with the WT ANXA2 plasmid restored radiation resistance (**Fig. 4E**, left well and bar graph). Together, these experiments show that Y23 phosphorylation is a critical modification that enables ANXA2 to bind miR-603, drive its IR-induced export through MV shedding, and confer radiation resistance.

### MV-mediated miR-603 export is driven by radiation-induced ROS

Because Y23 phosphorylation of ANXA2 is mediated by SRC^60,61^, and SRC is activated by reactive oxygen species (ROS)^62,63^, we hypothesized that IR-induced ROS serve as the upstream trigger for miR-603 release. To test this hypothesis, LN340 cells were subjected to IR or tert-butyl hydroperoxide 1 (t-BHP1, an oxidative stress-inducing agent), with or without N-acetylcysteine (NAC, an antioxidant), followed by assessment of MV and miR-603 release. DCF-DA flow cytometry revealed strong ROS induction by IR and t-BHP1, with NAC suppressing IR-induced ROS (**Fig. 5A**). Nanoparticle tracking analysis of EVs from IR-treated LN340 cells showed increased MV release (**Fig. 5B**, top right panel), an effect suppressed by NAC (**Fig. 5B**, bottom left panel). t-BHP1 treatment elicited a comparable increase in MV release relative to IR (**Fig. 5B**, middle right panel).

**Figure 5.**
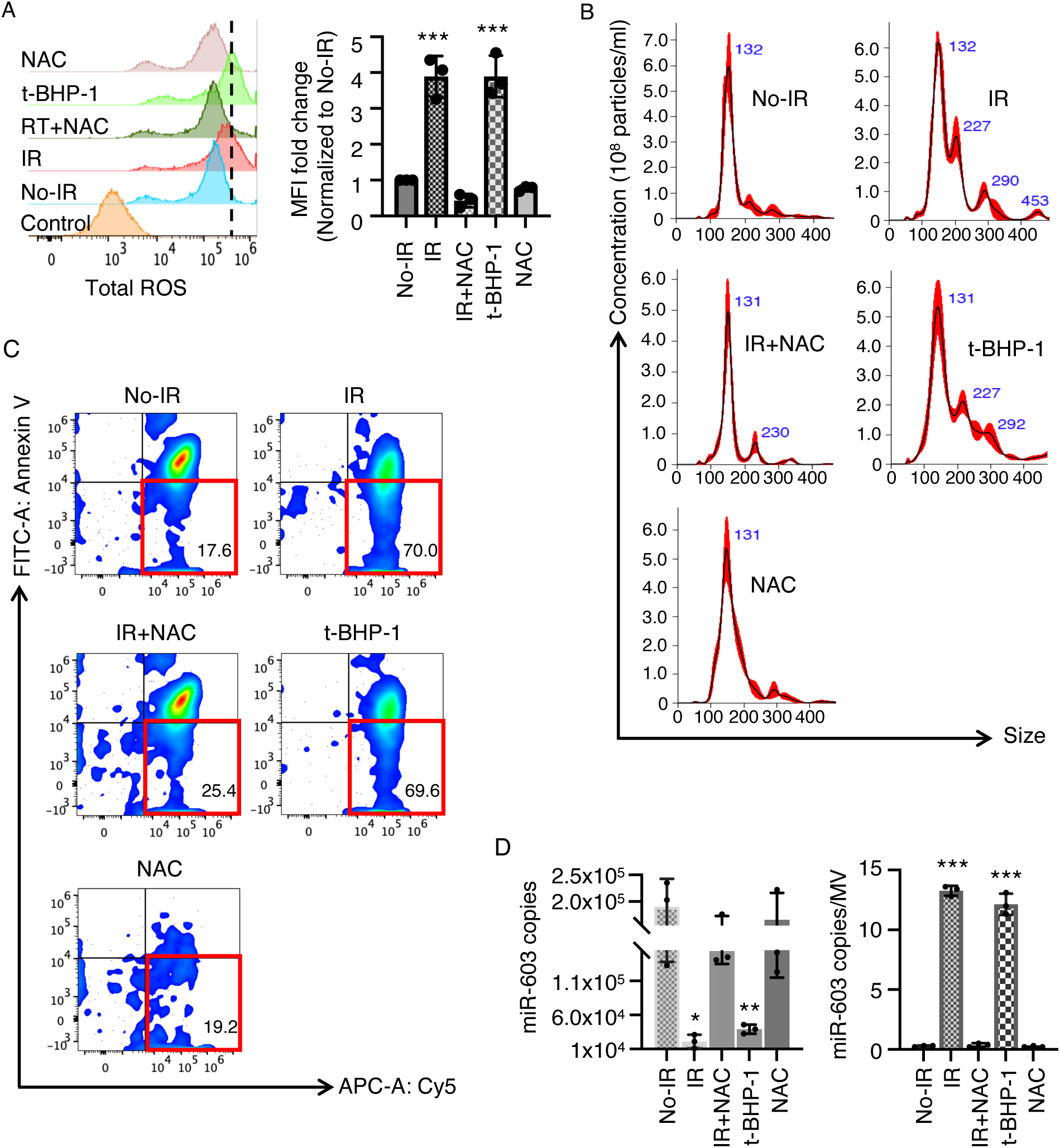
MV-mediated miR-603 export is driven by radiation-induced reactive oxygen species (ROS). **A.** Modulation of ROS by IR, tert-butyl hydroperoxide 1 (t-BHP1, to induce oxidative stress), and N-acetylcysteine (NAC, an antioxidant) treatment. LN340 cells were treated with DCF-DA, exposed to ionizing radiation IR (with or without NAC) or t-BHP1, and subjected to flow-cytometric analysis. DCF-DA fluorescence is shown in a histogram (left panel). Fold change in mean fluorescence intensity (MFI) is presented as a bar graph (right panel). **B.** ROS induced cellular release of MVs. EVs were collected from conditioned media from the above-treated cells and subjected to nanoparticle tracking analysis. Representative nanoparticle tracking shows the distribution of particle size as a function of particle number (particles/mL) and modal size (nm). **C.** ROS induced cellular release of miR-603 containing MVs. MVs were collected from conditioned media from the above-treated cells and subjected to Cy5/FITC-Annexin V flow cytometry. Results are shown in a contour plot. **D.** ROS induce shift of miR-603 from the cytoplasm to MVs. Total RNA was isolated from LN340 cells treated with IR, t-BHP1, NAC or combination of NAC or IR. MVs were isolated from the conditioned media, followed by miR-603 RT-qPCR. Data represent mean ± SD. *p< 0.05, **p< 0.01 and ***p< 0.001.

To determine whether the release MV contained miR-603, Cy5-labeled miR-603 into LN340 prior to treatment with IR, NAC, or RT+NAC. Flow-cytometric analysis of MVs from these cells showed that IR induced a ∼5-fold increase in Cy5⁺/Annexin V⁻ MVs (**Fig. 5C**, top right panel). This induction was abolished by NAC treatment (**Fig. 5C**, bottom left panel). Moreover, t-BHP1 treatment induced a level of Cy5⁺/Annexin V⁻ MVs comparable to that observed with IR (**Fig. 5C**, middle right panel). Finally, the IR-induced reduction in cytoplasmic miR-603 and corresponding increase in MV-associated miR-603 were suppressed by NAC treatment (**Fig. 5D**). And t-BHP1 exposure produced a reduction in cytoplasmic miR-603 and a corresponding rise in MV-associated miR-603 comparable to IR (**Fig. 5D**, right panel). Together, these findings support our hypothesis that ROS act as upstream signals that drive MV release and miR-603 export.

### ROS-dependent activation of SRC is responsible for ANXA2 Tyr23 phosphorylation

Since ROS activates SRC via sulfenylation of Cys185 and Cys277 (forming Cys-SOH)^41^, we next tested whether these SRC sulfenylation events are required for ANXA2 Y23 phosphorylation and miR-603 export. Consistent with published studies^64,65^, RT induced SRC activation, as evidenced by increased SRC Y416 phosphorylation and ANXA2 Y23 phosphorylation (**Fig. 6A**). To determine whether SRC activity is required for miR-603 export, cells were treated with a SRC inhibitor (A-419259 trihydrochloride), prior to IR. Twenty-four-hour treatment significantly reduced IR-induced increase of miR-603 in MVs, as quantified by RT-qPCR (**Fig. S12A**). These findings suggest that SRC signaling modulates miR-603 export through MVs.

**Figure 6.**
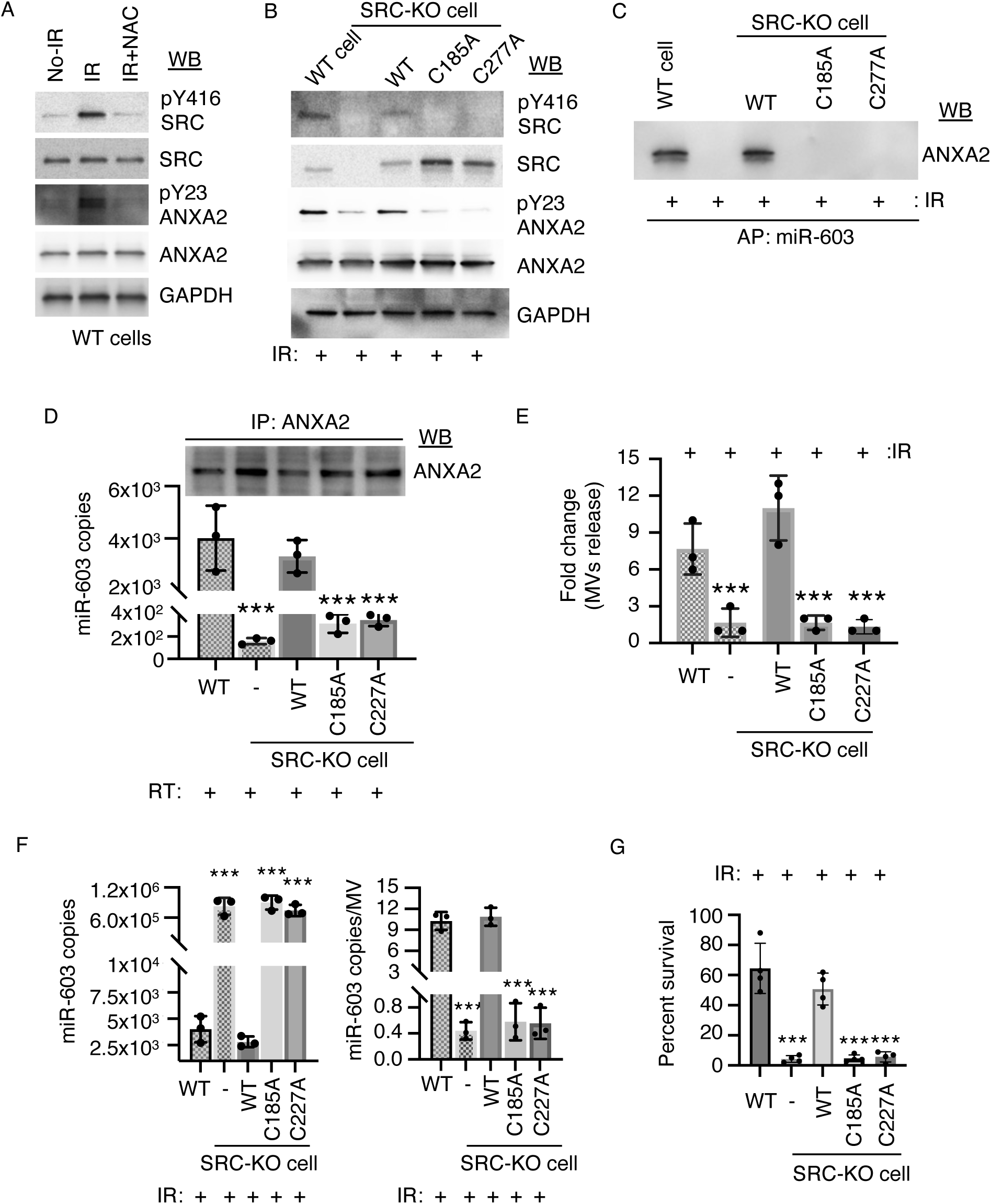
ROS-dependent activation of SRC is responsible for ANXA2 tyrosine 23 phosphorylation. **A.** ROS induced SRC Y416 phosphorylation. LN340 WT cells were treated with ionizing radiation, t-BHP1, NAC or combination of NAC and IR. The cell lysates were subjected to immunoblotting with indicated antibodies. **B.** SRC C185 and C277 required for SRC activation and ANXA2 Y23 phosphorylation. SRC KO LN340 cells were transfected with SRC WT or C185A and C277A mutant plasmids. The cell lysates were subjected to immunoblotting with indicated antibodies. **C.** SRC C185A or C277A mutations abolish ANXA2-miR-603 interaction. SRC-KO LN340 cells expressing WT, C185A, or C277A SRC were transfected with biotinylated miR-603. MVs isolated from conditioned media were subjected to streptavidin pulldown, and bound proteins were analyzed by ANXA2 immunoblotting. **D.** miR-603 is not enriched in ANXA2 immunoprecipitates from MVs of SRC C185A and C277A-expressing cells. MVs were isolated as described in (C) and subjected to ANXA2 immunoprecipitation, followed by RNA extraction and RT-qPCR analysis of miR-603. *<u>Top</u>*: ANXA2 immunoblot of immunoprecipitated samples. *<u>Bottom</u>*: miR-603 RT-qPCR of RNA isolated from the immunoprecipitates. Data represent mean ± SD. ***p< 0.001. **E.** SRC C185 and 277 is required for radiation-induced export of miR-603-containing MVs. Wild-type (WT) and SRC KO LN340 CD63-GFP cells were transfected with SRC C185A and 277A plasmid. Twenty-four hours post transfection, the cells were transfected with Cy5-miR-603, subjected to irradiation and monitored by live-cell imaging. Quantification of Cy5-miR-603^+^ MVs release (actual video shown in **Fig. S12B**) across 100 fields represented in bar graph. ***p < 0.001 between indicated groups (Student’s t-test). **F.** SRC C185A or C277A mutations abolish IR-induced redistribution of miR-603 from the cytoplasm to MVs. SRC KO cells were transfected with C185A or C277A mutants and treated with 6 Gy IR. Total RNA was isolated from the cells and corresponding MVs, followed by RT-qPCR quantification of miR-603. Data represent mean ± SD. ***p < 0.001. **G.** C185A and C277A mutants of SRC induces the radiation sensitivity. SRC KO LN340 cells were transfected with WT SRC or the Y23A mutant plasmids and subjected to clonogenic survival analysis. Representative images from the clonogenic survival assay are shown.

To define the role of SRC sulfenylation in miR-603 export, we generated CRISPR SRC KO cells and reintroduced WT SRC or the redox-insensitive alanine substitution C185A and C277A mutants. The IR-induced ANXA2 Y23 phosphorylation was greatly reduced in both the C185A and C277A mutants (**Fig. 6B**). Next, we tested whether these mutants support miR-603 binding to ANXA2. We transfected Bi-miR-603 into SRC KO LN340 cells expressing WT, C185A, or C277A SRC and then treated with IR. MVs were isolated, and the lysates were subjected to streptavidin affinity pulldown followed by ANXA2 immunoblotting. ANXA2 was detected in Bi-miR-603 pulldowns from cells expressing WT SRC. However, no detectable ANXA2 signal was observed in Bi-miR-603 pulldowns from cells expressing C185A or C277A mutants (**Fig. 6C**). Reciprocal immunoprecipitation using an anti-ANXA2 antibody was performed using the same MV lysates derived from cells expressing WT, C185A, or C277A SRC, followed by RNA extraction from the immunoprecipitated fractions and miR-603 RT-qPCR. This experiment revealed that IR induced an approximately an order-of-magnitude enrichment of miR-603 in immunoprecipitates derived from cells transfected with WT SRC. Such enrichment was not observed in immunoprecipitates from cells transfected with C185A or C277A SRC mutants (**Fig. 6D**).

Next, we tested whether SRC C185 and C277 sulfenylation are required for radiation-induced miR-603 export using live-cell imaging. This experiment showed that SRC KO cells reconstituted with the C185A or C277A mutants exhibited ∼5-fold reduction in MV-mediated release of Cy5-miR-603 relative to those reconstituted with WT SRC (**Fig. 6E, S12B (video)**). Finally, SRC-KO cells reconstituted with the C185A or C277A mutants also failed to show the IR-induced decrease in cytoplasmic miR-603 and corresponding increase in MV-associated miR-603 (**Fig. 6F**). These results suggest that redox-dependent SRC activity is essential for efficient MV-mediated export of miR-603 following irradiation.

Functionally, SRC KO cells exhibit a ∼ 5-fold increase in sensitivity to IR. This sensitivity was rescued by re-expression of WT SRC but not by the C185A or C277A mutants (**Fig. 6G**). Importantly, miR-603 transfection induced a comparable ∼5-fold increase in radiosensitivity in WT LN340 cells yet did not further enhance the IR sensitivity of SRC KO cells. This epistatic relationship supports a model in which SRC and miR-603 act within the same pathway governing radiation resistance.

## DISCUSSION

Beyond its well-known genotoxic effects^66–68^, IR induces cytoplasmic responses of comparable biological relevance^2^. A notable example involves the Ca²⁺-dependent recruitment of ANXA2 to the damaged plasma membrane, facilitating the stabilization of the injured surface and the shedding of damaged membranes through the release of ANXA2-containing MV^49^. Our findings support a model in which membrane repair is coupled to miRNA export to drive acquired radiation resistance. We found that IR-induced miR-603 export is mediated through MV release and requires ANXA2 binding to miR-603. Computational modeling suggests that phosphorylation of ANXA2 at Y23, a SRC phosphorylation site^69^, is required for this binding. Confirming this prediction, an ANXA2 Y23A mutant fails to bind miR-603. In SRC KO cell lines expressing SRC mutants (C185A and C277A) that cannot undergo ROS-mediated activation, both ANXA2-miR-603 binding and IR-induced miR-603 export were impaired. These findings support a model in which IR-generated ROS activate SRC through sulfenylation of Cys185 and Cys277, enabling ANXA2 Y23 phosphorylation. The phosphorylated ANXA2 binds miR-603, chaperons it to damaged membrane sites, where the ANXA2-miR-603 complex is subsequently released with the shed MVs (**Fig. 7**). While these observations were made in the context of glioblastoma, they point to a broader paradigm in which acute membrane stress drives selective RNA export through plasma-membrane–derived vesicles across diverse disease contexts.

**Figure 7.**
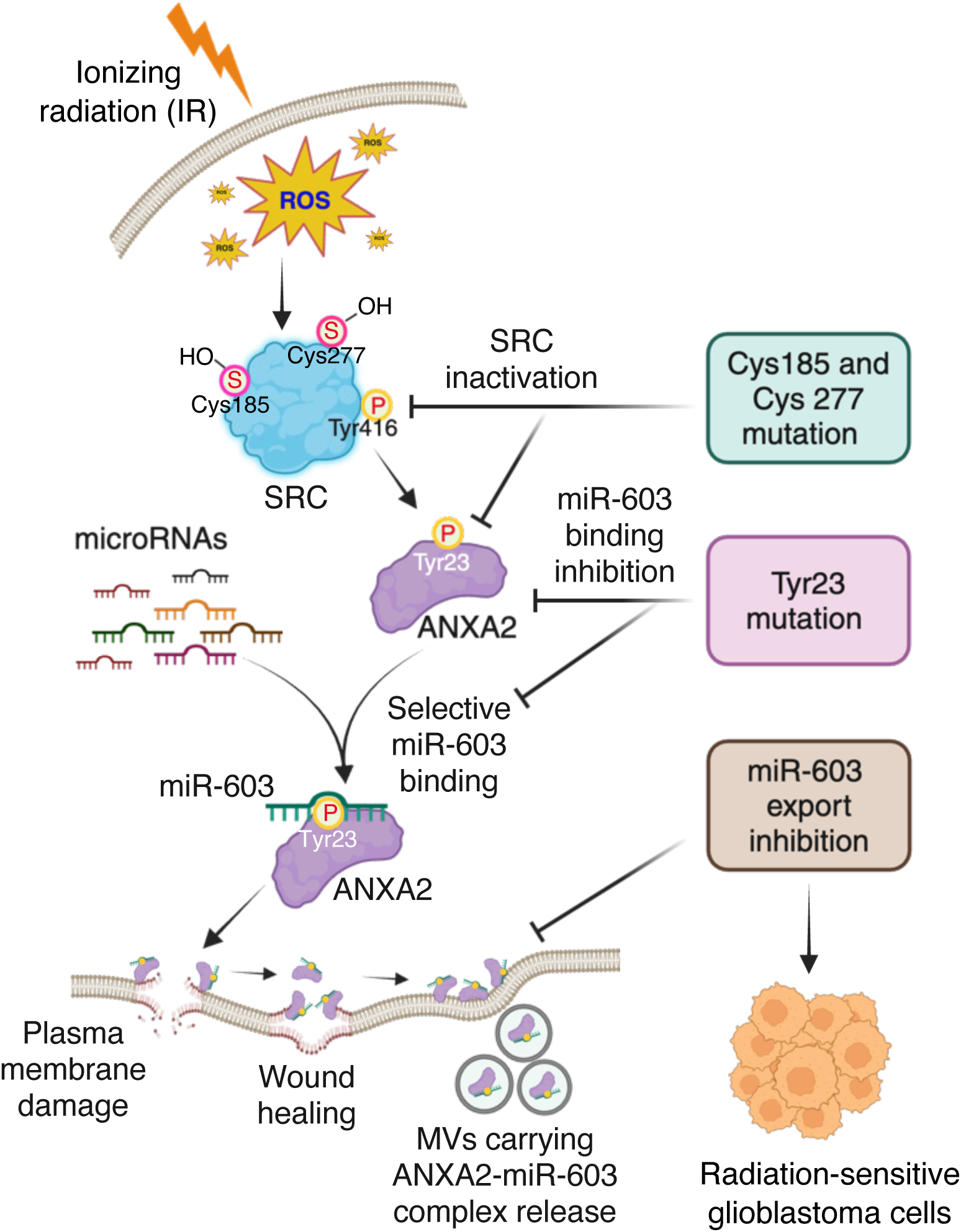
ROS-SRC-ANXA2 signaling drives selective microvesicle export of miR-603 in response to radiation. Ionizing radiation (IR) induces reactive oxygen species (ROS), which activate SRC kinase through redox-sensitive cysteine residues (Cys185 and Cys277). Activated SRC phosphorylates Annexin A2 (ANXA2) at Tyr23, enabling ANXA2 to selectively bind miR-603. The phosphorylated ANXA2-miR-603 complex is recruited to plasma membrane stress site, where membrane remodeling and wound-healing processes facilitate the formation of microvesicles (MVs). These MVs package and release the ANXA2-miR-603 complex into the extracellular space, resulting in intracellular depletion of miR-603. This ROS-SRC-ANXA2 axis represents a signaling circuit that couples radiation-induced membrane stress to selective miRNA export via extracellular vesicles. The figure created in BioRender.

SRC functions as a central node in cellular stress responses^70,71^, playing an essential role in survival-promoting signaling programs^72^. Although prior clinical studies of SRC inhibitors in glioblastoma have not yielded survival signals^73^, these trials largely targeted SRC as a constitutively active oncogenic driver rather than as a context-dependent mediator of therapy-induced adaptation^74,75^. Moreover, prior trials (NCT01331291, NCT00423735) were restricted to the recurrent setting and did not include combination with IR^76^. Our findings suggest that the biological complexity of SRC was not adequately considered in these clinical translation designs. Given the findings of this study and the broader literature, SRC inhibition should be evaluated in combination with radiation, a strategy supported by preclinical studies^77^. The availability of a clinical-grade, blood-brain barrier-penetrant SRC inhibitor potentially shortens the path to clinical translation^78^. Finally, our study indicates that SRC inhibition prevents radiation-induced release of miR-603, which compromises IGF signaling^9,79^. This strategy may be most effective in IGF-dependent cell states^80^, with cytoplasmic miR-603 levels serving as a potential biomarker for such states.

Targeting the SRC-ANXA2-miR603 axis additionally offers a potential strategy to reverse immune suppression in the glioblastoma microenvironment. Radiation-induced release of miR-603 reshapes the glioblastoma microenvironment through two convergent mechanisms. First, depletion of intracellular miR-603 in tumor cells derepresses IGF1 and IGF1R, amplifying IGF-axis signaling that drives the recruitment of immunosuppressive myeloid populations^81^. Second, the exported miR-603 containing MVs is taken up by the endothelial^82^ and immune cells in the tumor microenvironment^10^, and this uptake is enhanced by ANXA2-mediated interactions^82^. Within these cells, miR-603 suppresses IGF signaling. Reduced IGF-axis activity in microglia diminishes phagocytic capacity and promotes a shift toward a more tumor-permissive state^83,84^. In T cells, IGF suppression impairs metabolic fitness, proliferation, and effector cytokine production^85^. In other immune cells, including dendritic cells and macrophages, loss of IGF signaling weakens antigen presentation and blunts pro-inflammatory activation^86^. Together, these convergent effects likely contribute to an immunosuppressive niche that can potentially be reversed by modulating the SRC-ANXA2-miR-603 axis.

Computational modeling of the ANXA2 RNA-binding site suggests that miR-603 is unlikely to be the sole miRNA released during radiation-induced MV export; other miRNAs and possibly mRNAs ^8,87,88^ may be chaperoned by ANXA2 for MV release. Moreover, proteins shed alongside damaged membranes, whether binding partners of or independent of ANXA2, may themselves chaperone nucleic acids and proteins into MVs^89,90^. Systematic characterization of the full repertoire of proteins and molecules involved in this membrane repair-linked export pathway should provide critical insight into the broader biological impact of MV release as it relates to acquired radiation resistance. Such analysis has the potential to reveal a more global regulatory program in which membrane repair, RNA export, and protein scaffolding converge to shape cellular adaptation and microenvironmental signaling. This framework further highlights the broader tendency of cancers to redirect and exploit physiologic mechanisms, such as the membrane repair machinery, to adapt to evolutionary pressures imposed by therapeutic interventions.

From a systems perspective, biological processes operate under persistent energetic constraints^91^, with every cellular function competing against the entropic cost of maintaining order^92,93^. Within this energetic landscape, coupling multiple processes to a single energetic expenditure is favored. In other words, if plasma membrane repair and shedding of damaged membrane-containing MV are required for cellular survival, it is energetically efficient to couple this repair to the release of cargos that would promote cellular survival. The dual function of ANXA2 in orchestrating membrane repair and chaperoning miR-603 for export exemplifies this principle. This frugality is unlikely to be confined to membrane repair and microRNA export; rather, it likely reflects a broader organizing principle governing biological systems.

In summary, our findings suggest that IR activates a membrane-repair program in which SRC phosphorylates ANXA2, enabling its binding to miR-603. Recruitment of this complex to IR-damaged membranes and the incorporation of this complex into shedding vesicles result in the export of miR-603, contributing to radiation resistance. Clinically, these results position the SRC–ANXA2-miR-603 axis as a tractable therapeutic vulnerability.

## METHODS

### Cell culture

Human glioblastoma cell lines LN340, LN215, LN18, LN229 and SF767 as previously described^9,94^ and cultured in Dulbecco’s modified Eagle’s medium (DMEM) supplemented with 10% fetal bovine serum (FBS), 2 mM L-glutamine, and 1% penicillin-streptomycin. In parallel, patient derived glioblastoma lines GBM39, HK374 and MGG123 were grown as neurospheres in NeuroCult media (Gibco) supplemented with heparin, human epidermal growth factor (EGF), human fibroblast growth factor (FGF) and 10% fetal bovine serum (FBS) in ultra-low attachment flasks (Nunc). All glioblastoma lines used in this study were wtIDH/umMGMT and maintained in a humidified atmosphere at 37 ^◦^C incubator with 5% CO_2._ Cell line identity was verified annually by STR profiling, and cultures were routinely tested for mycoplasma contamination using Mycoplasma detection kit (ATCC).

### Extracellular vesicle (EV) isolation and quantification

LN340 cells were cultured in DMEM (Gibco) supplemented with 10% exosome-depleted FBS (Thermo Fisher Scientific) for 24 h. The culture medium was replaced prior to irradiation. Using a previously published protocol^9^, conditioned media collected 48 h post-irradiation and subjected to sequential centrifugation at 2,800 × g for 10 min (removing cells and debris) followed by intermediate speed at 10,000 × g for 1 h to sediment MVs. The resulting supernatant was then ultracentrifuged at 100,000 × g for 1 h to isolate exosomes. The MV and exosome pellets were washed, then resuspended in 500 µL 1× PBS (pH 7.4) containing 1 mM MgCl₂ and 25 U/mL Benzonase (Millipore Sigma), followed by incubation for 90 min at room temperature to remove nucleic acids bound to the outer membrane of vesicles. The samples were then diluted in 10 mL of 1× PBS and collected by centrifugation at 10,000 × g for MV, followed by ultracentrifugation at 100,000 × g for 1 h for exosomes.

EV size distribution and particle concentration were analyzed by nanoparticle tracking analysis using a NanoSight NS300 instrument equipped with NTA software 3.4 Build 3.4.4 (Malvern analytical). EV preparations were diluted in 1× PBS to obtain particle concentrations within the optimal detection range and analyzed using standardized acquisition settings.

### RNA isolation and quantitation

Total RNA or miRNA was isolated from cells or extracellular vesicles (EV) using the miRNeasy Kit (Qiagen) following the manufacturer’s instructions. cDNA was synthesized from the total RNA using Omniscript reverse transcription kit (Qiagen). Reverse transcription of miRNA was performed using the miRCURY LNA RT kit (Qiagen). mRNA and miR-603 transcripts were quantified by qPCR using target-specific primer sets or miRCURY LNA-enhanced primers (Qiagen: YP00204112) and SYBR Green (Bio-Rad) on a Bio-Rad Chromo4 DNA Engine real-time PCR system. To quantify the absolute copy number of miR-603, cycle threshold (Ct) values were calibrated using a standard curve derived from a synthetic human miR-603 mimic (Qiagen, MSY0003271) serially diluted to a final concentration of 1×10^1^, 1×10^2^, 1×10^3^, 1×10^4^, 1×10^5^, 1×10^6^, 1×10^7^, and 1×10^8^ copies. miRNA copies per EV were calculated by dividing the total EV numbers quantified by NanoSight.

### Plasmids and cell line generation

The full-length human ANXA2 coding sequence was amplified from cDNA generated using the Omniscript RT Kit (Qiagen) from total RNA isolated from LN340 cells. PCR amplification was performed using ANXA2-specific primers (Sup. Table 1) under the following conditions: initial denaturation at 94 °C for 2 min, followed by 30 cycles of denaturation at 94 °C for 30 sec, annealing at 52 °C for 30 sec, and extension at 72 °C for 1 min, with a final extension at 72 °C for 5 min. The amplified ANXA2 fragment was subsequently subcloned into the pCMV6-AC-GFP expression vector (OriGene) using MluI and XhoI restriction sites. ANXA2 S11A, Y23A, and S25A mutants were derived from the full-length ANXA2 using site-directed mutagenesis (SDM) primer pairs (Sup. Table 1) and the QuikChange Lightning SDM kit (Agilent) in accordance with the manufacturer’s instructions. pcDNA3.1+/c-(k)DYK plasmid containing full-length SRC was purchased from Genescript (OHU28514). SRC C185A and C277A mutants were derived from this plasmid using specific SDM primer pairs (Sup. Table 1) and QuikChange Lightning Site-Directed mutagenesis kit (Agilent). All clones were confirmed by DNA sequencing.

For genetic disruption, ANXA2 (Horizon Discovery) and SRC (BPS Biosciences) CRISPR lentivirus were purchased and transduced to LN340 cells at 1 multiplicity of infection (MOI). Individual clones were selected after two weeks selection in media containing 2 mg/ml puromycin (Gibco). Each clone was sequenced. Absence of ANXA2 and SRC expression was confirmed by RT-qPCR and Western blotting as previously described^94^.

### miRNA, siRNA and plasmid transfection

LN340 cells were transfected with 30 nM biotinylated miR-603 (Dharmacon), Cy5-labeled miR-603 (Dharmacon), control miRNA (Dharmacon), siANXA2 (Qiagen), or non-targeting siRNA (Qiagen) using Lipofectamine™ RNAiMAX (Invitrogen) according to the manufacturer’s instructions.

For genetic rescue or mutational analysis, LN340 ANXA2-KO cells were transfected with pCMV6-AC-GFP-ANXA2 or the indicated substitution mutants (S11A, Y23A, S25A). In parallel, LN340 SRC-KO cells were transfected with pcDNA3.1+/c-(k)DYK-SRC (GenScript) or cysteine substitution mutants (C185A, C277A), or with the corresponding empty vector control. All plasmid transfections were performed using X-tremeGENE HP DNA Transfection Reagent (Roche) according to the manufacturer’s protocol.

### Immunoprecipitation and Western blotting

For immunoprecipitation (IP) assays, 30 µL Dynabeads Protein G (Invitrogen-10003D) was washed twice with a 10-bead volume of IP lysis buffer (20 mM Tris-HCl (pH 7.4), 150 mM NaCl, 2 mM EDTA, and 1% NP-40), resuspended in IP lysis buffer with 1 mM BSA, and incubated with anti-mouse ANXA2 antibody (Santa Cruz Biotechnology; sc47696), anti-mouse M2-FLAG antibody (Sigma), or anti-mouse IgG control (Santa Cruz Biotechnology; sc-2025) for 45 min at room temperature. Bead-antibody complexes were washed once with IP wash buffer and incubated with 300 µg of total cell lysate at 4°C for 2 h with end-over-end rotation. Following incubation, immune complexes were washed four times with IP wash buffer, and bound proteins were eluted by boiling in 1× SDS sample buffer.

For Western blotting, the lysates or protein mixtures were subjected to 4-20% gradient SDS/PAGE electrophoresis, followed by transfer to nitrocellulose membrane (Biorad). Membranes were blocked with 5% BSA for 1 h at room temperature, then incubated overnight at 4◦C with primary antibody. After washing with 1x TBST buffer, membranes were incubated with horseradish peroxidase-conjugated secondary antibodies (Cell Signaling Technology) for 1 h at room temperature and developed using chemiluminescence reagent (GE biosciences) and imaged using the Bio-Rad ChemiDoc MP Imaging System.

### Flowcytometric analysis

For experiments involving analysis of EV Cy5-miR-603, LN340 cells were transfected with Cy5-miR-603 or miR-ctrl and then subjected to radiation or mock treatment. EVs were collected as described above and resuspended in 200 µL Annexin-V binding buffer (Thermo Scientific), stained with FITC-conjugated Annexin-V (Biolegend-640945), washed and resuspended with 1X FACS buffer, and analyzed using the BD FACSCalibur flow cytometer (BD Biosciences). DCFH-DA (Sigma-Aldrich) and OxiVision™ Blue Peroxide Sensor assays (AAT Bioquest) were performed following the manufacturer’s instructions. Flow cytometric analyses were performed using FlowJo software.

### Clonogenic and comet assays

For clonogenic assays, cells exposed to IR (6 Gy) or mock treatment were seeded into six-well plates at a density of 500 or 1,000 cells per well and cultured for approximately 14-21 days to allow colony formation. Colonies were fixed with 100% methanol for 30 min (room temperature), air-dried for an hour, and stained with crystal violet (0.5% in methanol) for 30 min. Excess stain was removed by repeated washing with deionized water, and plates were air-dried at room temperature overnight. Plate images were acquired with a Leica DMi8 microscope and quantified using LAS X (Leica Microsystems). The comet assay was performed using the Comet Assay Kit (3-well slides) (Abchem) according to a previously published protocol^9^, imaged using the Leica-DMi8 fluorescence microscope, and analyzed using the LASX application suite.

### Live-cell imaging

A CD63-GFP-expressing LN340 stable cell line was generated by transducing LN340 cells with CMV-CD63-GFP lentivirus (Creative Biogene). Stable clones were selected using puromycin (2 mg/mL). For imaging experiments, approximately 500 LN340 cells stably expressing CD63-GFP were seeded into four-compartment 35-mm polymer coverslip-bottom dishes (ibidi). Cells were transfected with Cy5-labeled miR-603 or Cy5-control miRNA, followed by 6 Gy IR. After treatment, cells were washed and replenished with FluoroBrite DMEM (Gibco) for live-cell imaging.

Time-lapse imaging was performed using a Nikon Ti microscope equipped with a Plan Apo 40× DIC M N2 air objective (NA = 0.95) within a Nikon A1Rsi system operating in widefield mode. Fluorescence excitation was provided by a Sola LED illuminator using Chroma 4906 (590-650 nm) and Chroma 49002 (450-490 nm) filter cubes, with emission detection at 662-737 nm (Cy5) and 500-550 nm (GFP). Sixteen-bit images were captured every 5 min for 48 h using an ORCA Flash 4.0 CMOS camera (Hamamatsu). With 1.5× zoom, brightfield images were acquired with 10 ms exposure (12 V illumination), while Cy5 and GFP fluorescence images were captured with 100 ms and 60 ms exposure times, respectively. Focus stability was maintained using the Perfect Focus System coupled with an MCL NanoDrive Piezo Z stage. Live cells were supported by a Tokai Hit environmental system (Tokai Hit, Gendoji-cho, Fujinomiya-shi, Shizuoka-ken, Japan) that maintained a humidified atmosphere with 5% CO_2_ at 37°C.

For imaging analysis, a General Analysis 3 routine was constructed in Nikon Elements Analysis software, version 5.42 (Nikon). Background was removed from the far-red and green fluorescence channels using rolling-ball subtraction with radii of 31 pixels (3.358 µm) for the far-red channel and 60 pixels (6.500 µm) for the green channel. Autoshading correction was applied to the brightfield channel using the Average Background setting. To correct bleaching, the Equalize Intensity function was applied to both fluorescence channels using the first-frame histogram as the reference. Local contrast was then enhanced to 75% with an 85 µm radius. For the brightfield channel, auto-contrast was applied with low and high thresholds of 0.005% and 0.009%. Cells were segmented using the Homogeneous Area Detection (edge-based) function with a compactness of 4.87 and a threshold of –7. Green-fluorescent vesicles were detected using an intensity threshold of 6,044 with a 0.5 µm minimum size, followed by 0.11 µm cleaning and 0.11 µm separation. Red vesicles were thresholded at 7,000 with 0.11 µm smoothing, 0.11 µm cleaning, 0.65 µm separation, and a 1.00 µm minimum size. A Having operation was then applied between the segmented extracellular space and the thresholded green- or red-fluorescent EVs, and vesicles were counted and measured within the GA3 routine.

### RNAscope *in situ* hybridization

Formalin-fixed, paraffin-embedded (FFPE) cell pellets were prepared following the Advanced Cell Diagnostics (ACD) RNAscope® technical guidelines with minor modifications. In brief, 2 × 10⁷ LN340 cells were treated with 6 Gy IR, collected by a 10 min centrifugation spin at 250 × g, washed with 1× PBS, and resuspended in 10 mL of freshly prepared 10% neutral buffered formalin. Samples were incubated at room temperature for 24 h on a rotator, washed, and resuspended in 200 µL of warm Histogel (epredia-HG-400-012-10 mL). Excess Histogel was removed by centrifugation (time, speed), and the pellet was allowed to solidify for 2-3 min on ice. Solidified Histogel cell pellets were submerged in 1× PBS, dehydrated, and embedded in paraffin. The blocks were hybridized with miR-603-specific or control miRNA probes, followed by signal amplification and chromogenic detection. Nuclei were counterstained with hematoxylin. Intracellular and extracellular miR-603 puncta were quantified from 12 fields.

### Transmission electron microscopy (TEM)

MV and exosome pellets were resuspended in 15 µL of 1× PBS (pH 7.4) and fixed by adding an equal volume of 4% paraformaldehyde. EVs were adsorbed onto glow discharged carbon-coated TEM grids by placing the grids onto 5 µL droplets of the EV suspension and incubated for 1 min. The excess samples were blotted with a filter paper. Grids were then gently washed with water and subjected to negative staining with 2% uranyl acetate for 30 sec at room temperature. Following staining, grids were embedded in 2.5% methylcellulose to enhance structural preservation and allowed air dry for 10 min prior to imaging.

Electron micrographs were acquired using a Hitachi HT7800 transmission electron microscope (Tokyo, Japan) equipped with a MegaView 3 digital camera and Radius 2.0 imaging software (EMSIS, Germany). Vesicle size measurements were performed on 15 randomly selected micrographs per condition using the arbitrary line measurement tool in Radius 2.0. EV size distributions were visualized using frequency histograms and scatter plots, where individual vesicle diameters were plotted with corresponding median values and ranges.

### Proteomic identification of miR-603 binding proteins

Three hundred microgram of EV proteins isolated from LN340 cells transfected with biotinylated-miR-603 (Bi-miR-603) or Bi-miR-Ctrl. The cells were treated with 6 Gy IR or mock followed by MV isolation from the conditioned media. The MVs were lysed by IP lysis buffer (20 mM Tris-HCl [pH 8.5], 150 mM NaCl, 2 mM EDTA, 1% Nonidet P-40) and 300 μg of MV protein was incubated with pre-equilibrated 25 µL Dynabeads M-280 Streptavidin beads (Invitrogen) at 4°C overnight with continuous rotation. The beads were collected by centrifugation, washed 4 times with IP wash buffer (20 mM Tris-HCl [pH 7.4], 300 mM NaCl, 0.5% NP-40) and subjected to competitive biotin elution in 250 μL of elution buffer containing 20 mM Tris-HCl (pH 8.5), 150 mM NaCl, 2 mM EDTA, 1% Nonidet P-40, 50 U RNaseOUT, and 4 mg/mL D-biotin^95^. The suspension was incubated at room temperature for 60 min with gentle agitation to prevent bead sedimentation. Following incubation, samples were centrifuged at 200 × g for 1 min, and the supernatant containing Bi-miR-603-associated complexes was collected. The elution step was repeated three times, and the eluates were lysed in SDS-sample buffer and subjected to 4-20% gradient SDS/PAGE electrophoresis, followed by silver staining (Bio-Rad). The gel bands were collected and processed for in-gel digestion^96,97^.

Peptides from the digestion were reconstituted in H₂O:acetonitrile:formic acid solution at a ratio of 97.9:2.0:0.1. Each sample (300 ng) was analyzed by capillary LC-MS/MS (Thermo Scientific) Dionex UltiMate 3000 RSLCnano system coupled to an Orbitrap Eclipse mass spectrometer (Thermo Scientific) equipped with FAIMS separation. Peptides were directly injected into load solvent and separated on a self-packed C18 column (ReproSil-PUR C18aq, 1.9 µm, 120 Å; 100 µm × 40 cm) at 55 °C. Chromatography was performed using solvent A (0.1% formic acid in H₂O) and solvent B (0.1% formic acid in acetonitrile) with the following gradient: 5% B (0–2 min), 8% B (2.5 min), 21% B (50 min), 32% B (75 min), and 90% B (85 min) with a flow rate of 350 nL/min (0–2.5 min) and 315 nL/min (2.5–85 min). FAIMS settings were: nitrogen cooling gas 5.0 L/min, carrier gas 4.6 L/min, electrode temperatures 100 °C, and CVs of −45, −60, and −75 V (1 s each).

MS1 spectra were acquired in the Orbitrap (Thermo Scientific) at 120,000 resolutions (380–1400 m/z), 50 ms injection time, and 100% automatic gain control (4 × 10⁵). Data-dependent MS2 scans were triggered for precursors with 2–5 charges and >2.5 × 10⁴ intensity using high-energy collisional dissociation fragmentation. MS2 settings were: 1.2 Da isolation window, 30% collision energy, Orbitrap detection at 30,000 resolution, first mass 110 m/z, 54 ms maximum injection time, and AGC (5×10⁴). Dynamic exclusion was set to 10 sec with ±10 ppm mass tolerance; exclusion lists were not shared among CVs. ESI voltage was +2.1 kV and the ion transfer tube was held at 275 °C.

Spectra were processed using SEQUEST (Thermo Scientific) in Proteome Discoverer 2.5. Searches were performed against the human UniProt database (UP000005640; downloaded Sept 20, 2021), merged with the common contaminant proteins database (http://www.thegpm.org/cRAP/index.html). Mass recalibration was applied (20 ppm precursor tolerance, 0.06 Da fragment tolerance) with fixed carbamidomethylation (CAM) modification of cysteine (57.0215 *m/z*). SEQUEST database search parameters included: trypsin specificity with one missed cleavage site, 15 ppm precursor tolerance, and 0.06 Da fragment ion tolerance. Static modification: CAM-Cys (+57.021 Da). Dynamic modifications: protein N-terminal acetylation (+42.011 Da), Met oxidation (+15.995 Da), Gln-to-pyro-Glu (−17.027 Da), N-terminal Met loss (−131.040 Da), N-terminal Met loss + acetylation (−89.030 Da), and Asn/Gln deamidation (+0.984 Da). We applied 1% protein and peptide False Discovery Rate (FDR) filters using the Percolator algorithm (https://doi.org/10.1038/nmeth1113) in the protein database.

### *In silico* mutagenesis of ANXA2 structural modeling

Crystal structures of ANXA2 were obtained from the RCSB Protein Data Bank (http://www.rcsb.org), including the wild-type protein (PDB ID: 5LPU) and phosphorylated ANXA2 (pANXA2; PDB ID: 5LQ0). Substitution mutation was introduced at Tyr23 of ANXA2 to alanine (Y23A) using Maestro (Schrödinger Release 2023). For all simulations, chains C and D were removed, retaining only chains A and B. Structural optimization of wild-type, phosphorylated, and mutant ANXA2 was performed using Avogadro with the MMFF94s force field and steepest-descent algorithm until convergence (ΔE ≈ 0).

The three-dimensional structure of miR-603 was modelled using the RNAComposer web server (https://rnacomposer.cs.put.poznan.pl/), an automated RNA tertiary structure prediction platform based on the machine-translation principle. RNA Composer generates 3D RNA structures by mapping secondary structure elements onto experimentally validated motifs curated in the FRABASE database. During model generation, the following parameters were applied: generation of A-form RNA-based double helices and single strands was disabled; only X-ray-determined structures were used as templates; and the resolution threshold was set to 3.0 Å. The resulting RNA 3D models were downloaded into Protein Data Bank (PDB) format and used for downstream docking and molecular dynamics analysis.

Details of molecular docking, dynamic simulations, and binding free-energy calculations are described in the Supplementary Information.

### Statistical analysis

Three or more independent experiments were performed for each assay and results were combined to define as mean ± SD. Statistical analysis were conducted using Microsoft Excel and GraphPad Prism software 10. The statistical significance was evaluated using an unpaired two-tailed Student’s t-test or one-way ANOVA. Following ANOVA, post hoc multiple comparisons were conducted using Tukey’s or Dunnett’s or Mann-Whitney test to identify statistically significant differences between specific group pairs. p value (*) of ≤0.05 was considered statistically significant.

## Acknowledgments

We thank Ananta Arukha for valuable technical assistance and for performing key preliminary experiments that contributed to this study. We are grateful to Dr. Amar Singh (Department of Surgery, University of Minnesota) for insightful discussions and guidance on extracellular vesicle (EV) flow cytometry analysis. We acknowledge Mary E. Brown and the University Imaging Centers (UIC), University of Minnesota, for their technical support with live-cell imaging and EV analysis. We also thank the Center for Metabolomics and Proteomics (CMSP), University of Minnesota, for assistance with EV proteomics. Transmission electron microscopy work was carried out in the Characterization Facility, University of Minnesota, which receives partial support from the NSF through the MRSEC (Award Number DMR-2011401) and the NNCI (Award Number ECCS-2025124) programs. This work was supported by start-up funding from the Department of Neurosurgery, University of Minnesota Medical School–Twin Cities awarded to G.S.

## Author Contributions Statement

G.S. and C.C.C. conceived and designed the study. G.S. supervised the research. S.S., I.M. and G.S. performed the experiments. A.K.M. performed bioinformatic analysis of molecular docking and molecular dynamics simulations. S.K. and A.R.N. contributed new reagents/analytic tools. S.S., A.K.M., I.M., S.K., A.R.N., C.C.C. and G.S. analyzed data. S.S., C.C.C. and G.S. drafted and critically revised the manuscript.

