## Supplemental Information for "A SRC-Annexin A2 axis that couples membrane repair to microRNA export during radiation stress in glioblastoma"

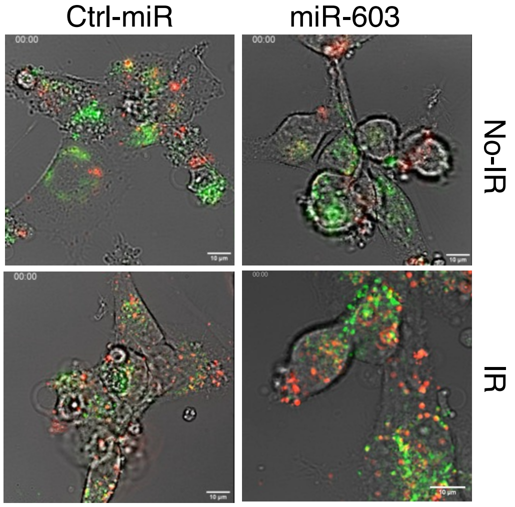


**Figure S1. Radiation-induced release of microvesicles containing miR-603.**

LN340 cells stably expressing CD63-GFP transfected with Cy5 tagged miR-603 or miR-NT (Ctrl-miR). The cells were treated with 6 Gy ionizing radiation (IR) or no-IR (0 Gy), subjected to live-cell imaging. Representative live-cell videos are presented. *Top left*: Ctrl-miR (0 Gy); *bottom left*: Ctrl-miR (6 Gy). *Top right*: miR-603 (0 Gy); *bottom right:* miR-603 (6 Gy). Scale bar is 10 µm.

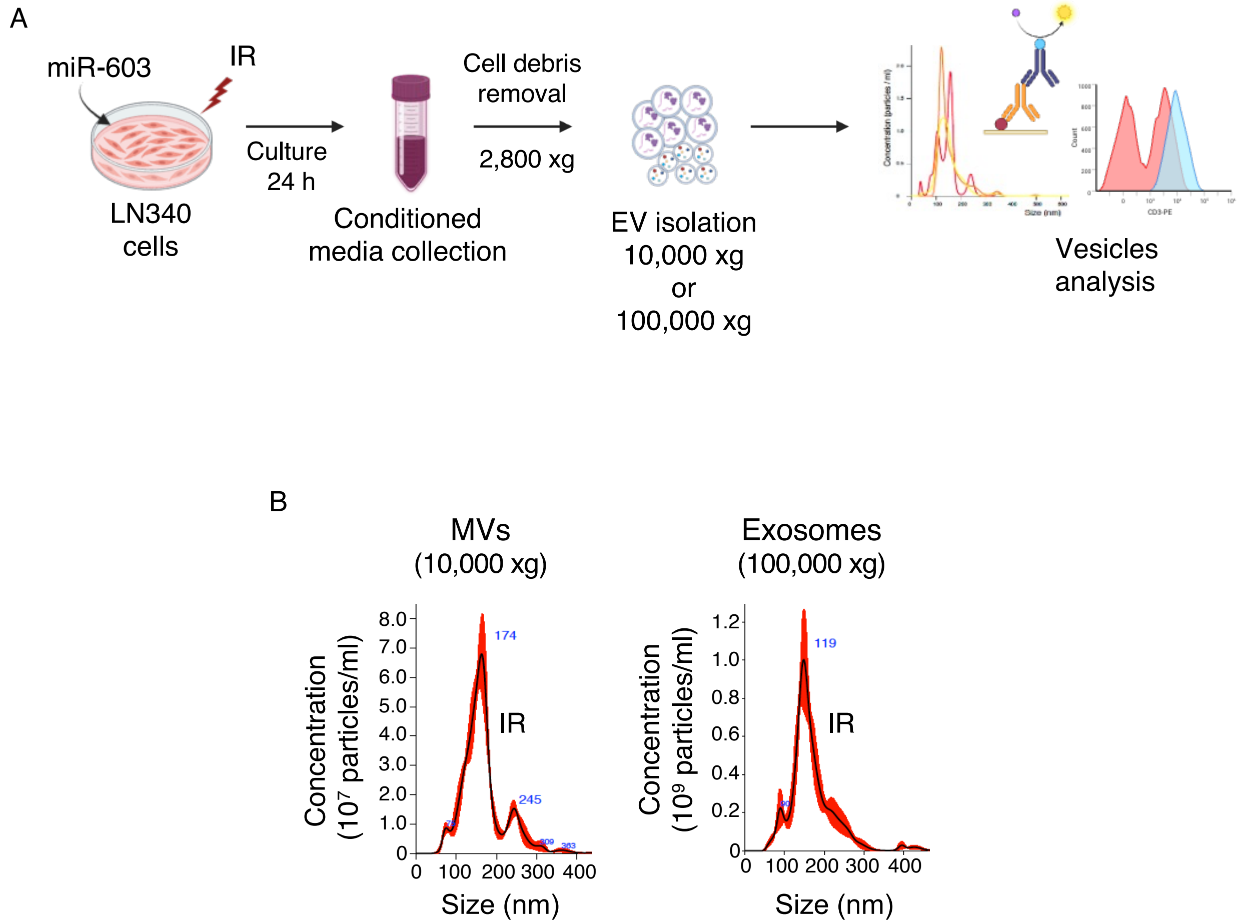


**Figure S2. Isolation and characterization of extracellular vesicle subtypes by differential centrifugation.**

**A.** Schematic representation of extracellular vesicle (EV) isolation from conditioned media using differential centrifugation, enabling separation of microvesicles (MVs) (10,000 × g) and exosomes (100,000 × g). **B.** Nanoparticle tracking analysis (NTA) of MVs and exosomes isolated by differential centrifugation. Representative NTA traces illustrate particle size distribution, and quantification shows total particle concentration (particles/mL) and modal particle size (nm).

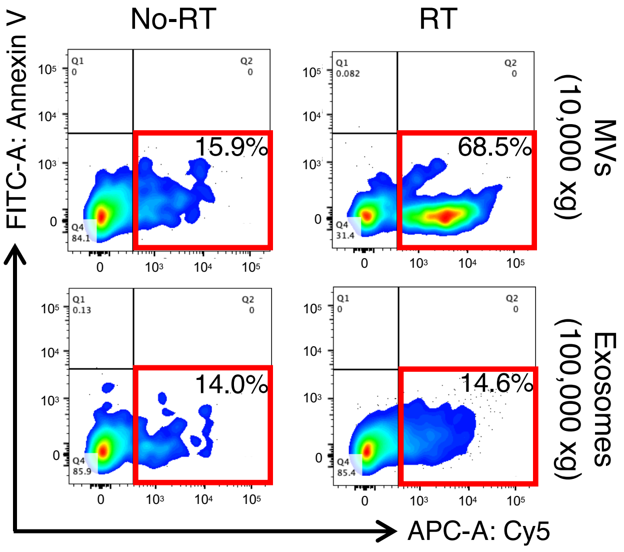


**Figure S3. Radiation induced microvesicle mediated miR-603 export.**

Differential ultracentrifugation-based isolation of microvesicles (MVs) (10,000 × g) and exosomes (100,000 × g) from media depleted of cellular debris. MVs and exosomes were isolated from Cy5-miR-603 transfected LN340 cells (with or without radiation treatment) and subjected to Cy5/FITC-Annexin V flow cytometry. Red box represents the $Cy5^{+}$-miR-603/FITC-Annexin $V^{-}$ MVs.

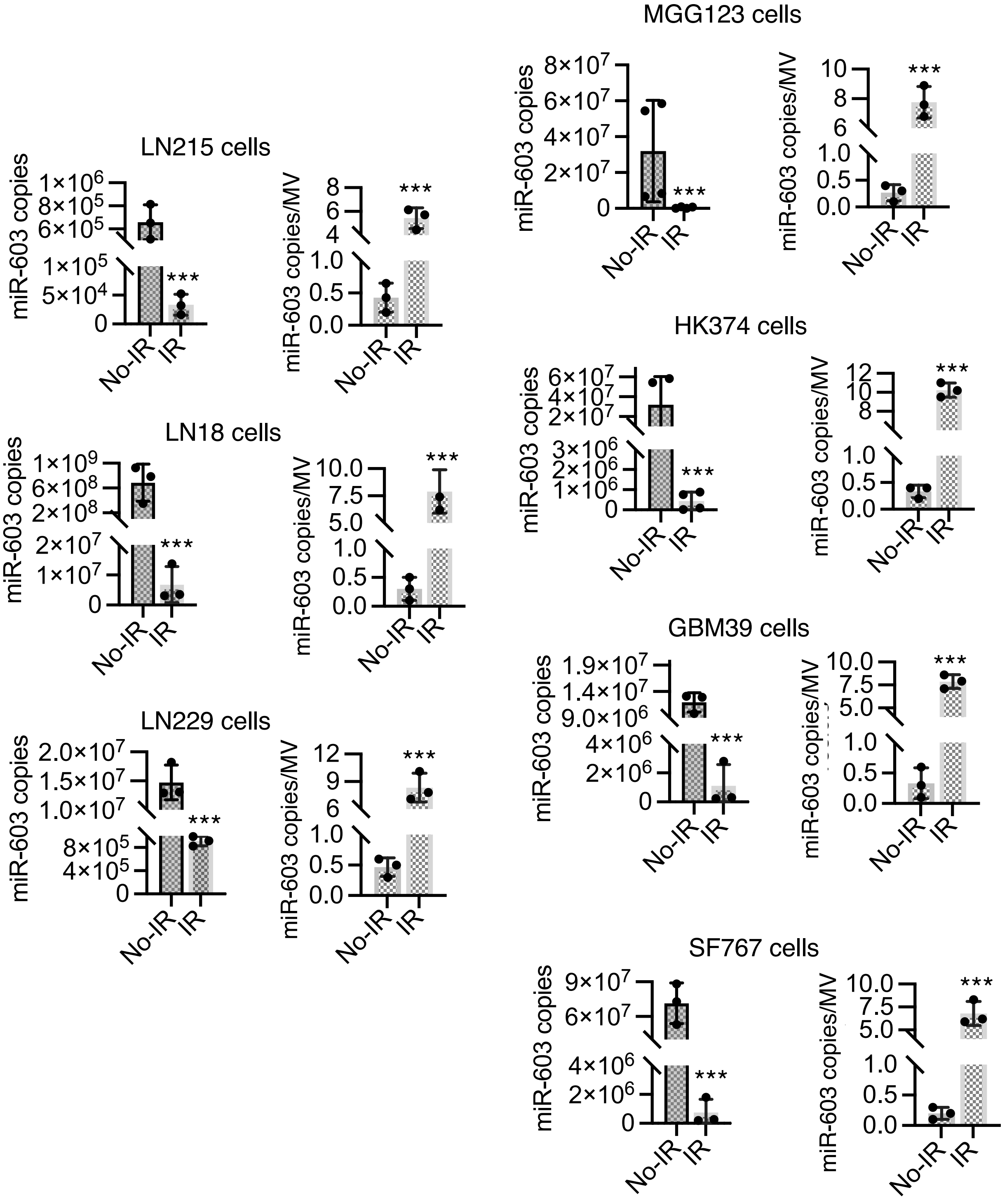


**Figure S4. Radiation induced microvesicle mediated miR-603 export.**

miR-603 levels in cytoplasmic and MV fractions. Indicated glioblastoma cells were treated with 6 Gy IR or mock control. RNA was extracted from cells as well as from MVs, and miR-603 levels were quantified by RT-qPCR.

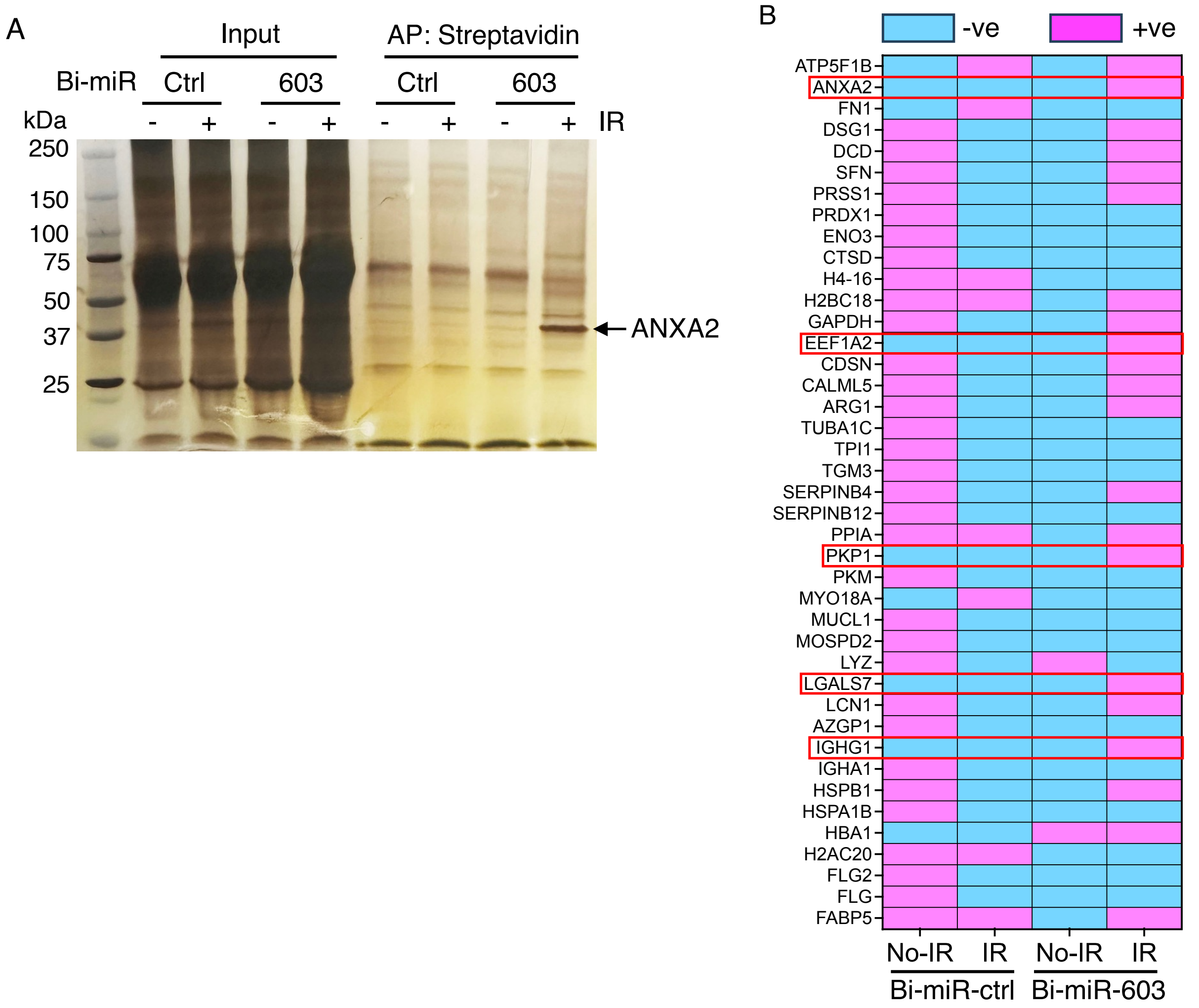


**Figure S5. Ionizing radiation promotes miR-603 association with Annexin A2 and microvesicle release.**

Streptavidin affinity pulldown of biotinylated miR-603 and candidate protein identification. LN340 cells transfected with biotinylated miR-603 (Bi-miR-603) or control miRNA (Ctrl-miR) and treated with ionizing radiation (IR). Microvesicles (MVs) isolated from the condition media were lysed and subjected to streptavidin pulldown. **A.** Silver staining identified enrichment of specific protein with miR-603 IR at ~ 39 kDa and that identified as Annexin A2 by mass spectrometry. **B.** Candidate proteins that interact with biotinylated-miR-603 (Bi-miR-603) in a radiation-dependent manner. MVs were isolated from irradiated and non-irradiated LN340 cells transfected with biotinylated-miR-603 (Bi-miR-603) or a biotinylated-control-miRNA (Bi-ctrl). The biotinylated RNAs were affinity-purified, and associated proteins were subsequently eluted and subjected to proteomic profiling. Candidates identified under each of the four conditions are shown in tabular format. Red indicates presence, and blue indicates absence. The red box indicates candidates found only in biotin-miR-603-pulled-down samples from irradiated LN340.

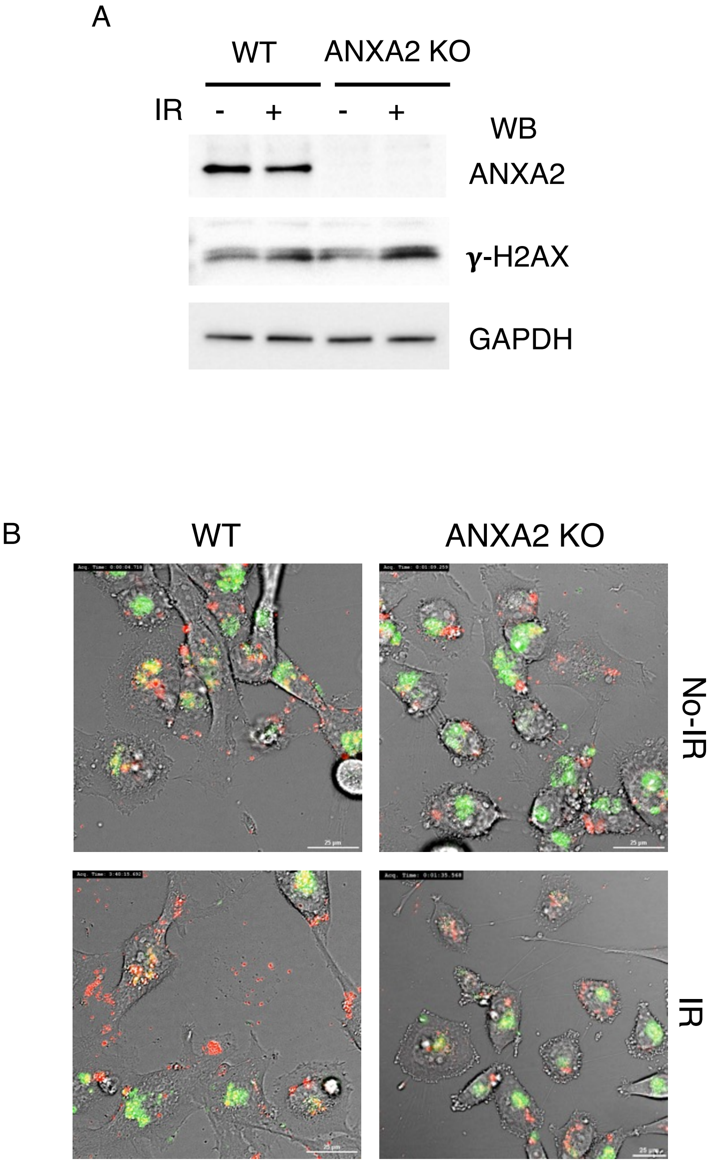


**Figure S6. ANXA2 knockout abolishes microvesicle mediated miR-603.**

**A.** LN340 WT and ANXA2 KO cells were treated with 6 Gy or 0 Gy ionizing radiation (IR). The cell lysates were analyzed by immunoblotting of indicated antibodies. **B.** LN340 WT and ANXA2 KO cells stably expressing CD63-GFP transfected with Cy5 tagged miR-603. The cells were treated with 6 Gy ionizing radiation (IR) or no-IR (0 Gy), subjected to live-cell imaging. Representative live-cell videos are presented. *Top left*: WT cells (0 Gy); *bottom left*: WT cells (6 Gy). *Top right*: ANXA2 KO (0 Gy); *bottom right:* ANXA2 KO (6 Gy). Scale bar is 10 µm.

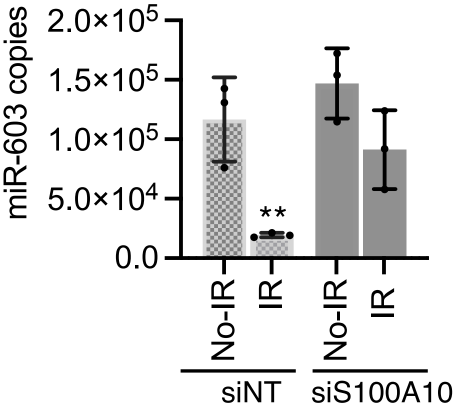


**Figure S7. Downregulation of S100A10 affects miR-603 microvesicle mediated export.**

Small interfering RNA (siRNA) targeting S100A10 was transfected into LN340 cells followed by ionizing radiation (IR) or mock treatment. Total RNA was isolated from siS100A10-transfected cells and from the corresponding microvesicles (MVs), followed by RT-qPCR quantification of miR-603. Data represent mean ± SD. **p < 0.01.

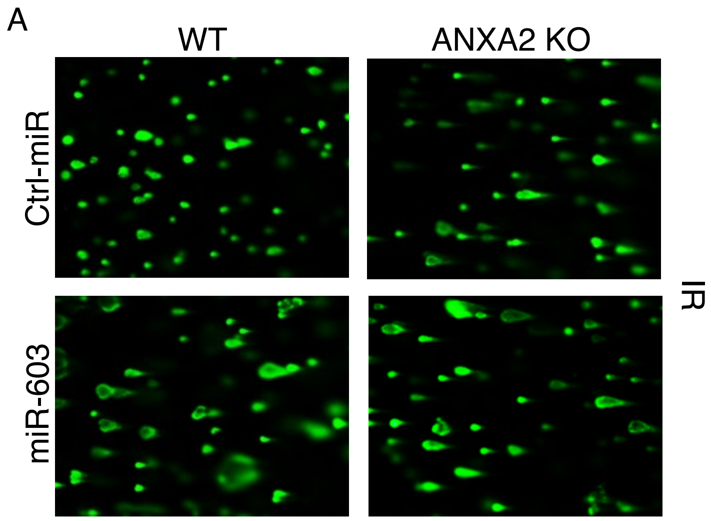


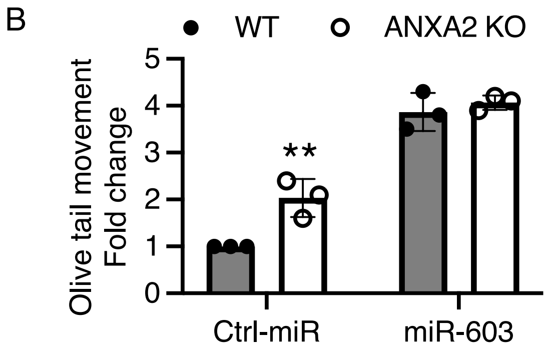


**Figure S8. ANXA2 and miR-603 required for ionizing radiation sensitivity.**

**A.** miR-603 increases comet tail moment in response to IR. LN340 WT and ANXA2 KO cells transfected with miR-603 or ctrl-miR and treated with IR. Comet assay was performed and the representative images of olive moment. Scale bar is 25 µm. **B.** Quantification of comet tail moment in irradiated LN340 WT and ANXA2 KO cells transfected with miR-603 or ctrl-miR. **$p < 0.01$ between indicated groups (Student's *t*-test).

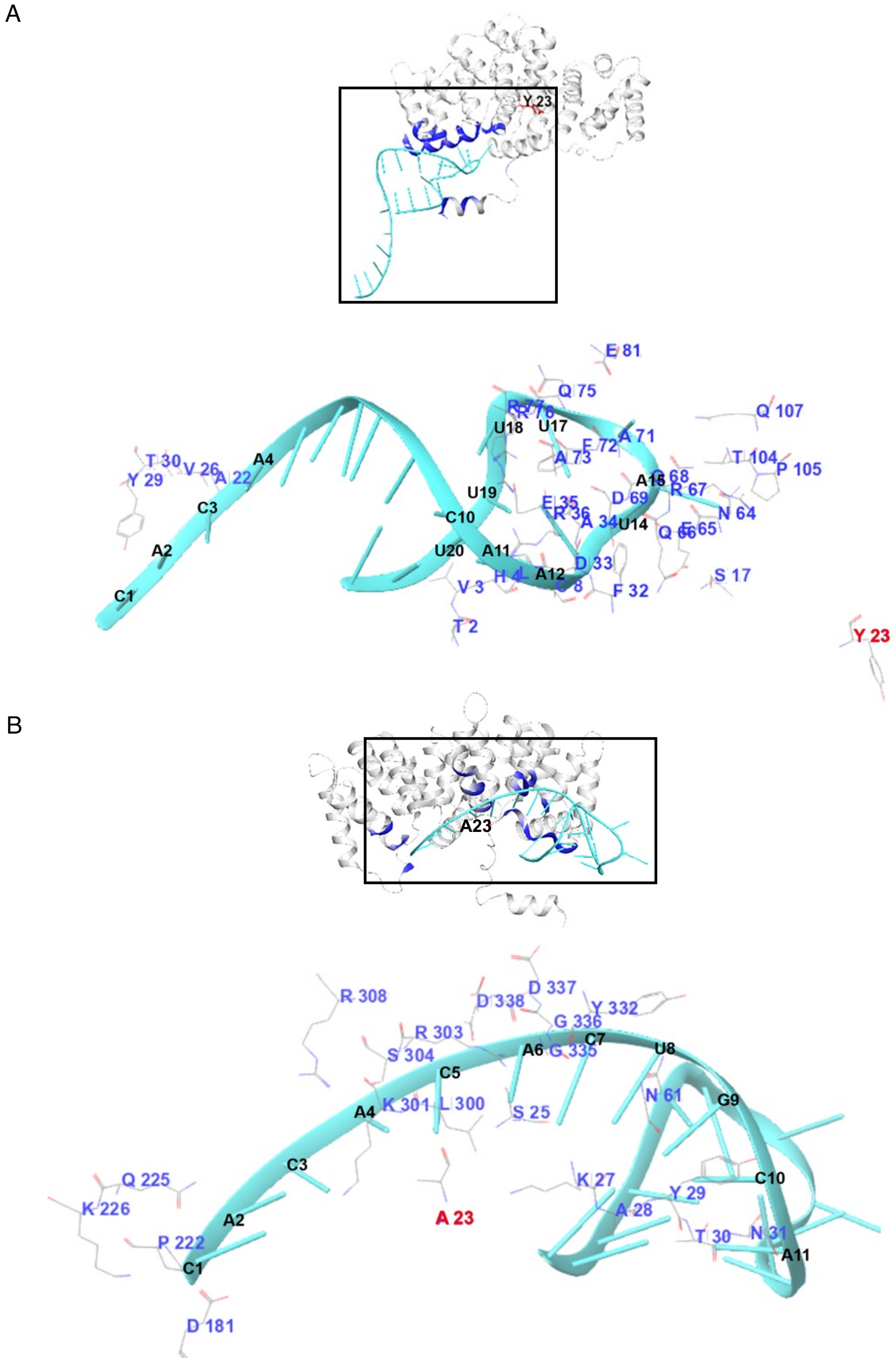
A
Y 23
E 81
Q 75
Q 107
U18
U17
A 71
F 72
A 73
T 104
P 105
T 30
V 26
A4
A 22
Y 29
U19
A15
D 69
R 67
C3
C10
E 35
R 36
N 64
A 34
U14
Q 66
U20
A11
D 33
A2
A12
S 17
V 3
F 32
C1
T 2
Y 23
B
A23
R 308
D 337
D 338
Y 332
R 303
G 336
C7
U8
A6
G 335
S 304
C5
N 61
G9
A4
K 301
L 300
S 25
C3
C10
Q 225
K 27
Y 29
K 226
A 28
A2
A 23
P 222
T 30
N 31
A11
C1
D 181

**Figure S9. Molecular Dynamics (MD) simulation of miR-603 with ANXA2 and ANXA2-Y23A.**

ANXA2: gray and blue; miR-603: cyan ribbon; miR-603 interacting region of ANXA2 shown in blue. ANXA2 residues: red/gray stick. miR-603 nucleotides shown in black. Y23 and mutant Y23 (A23) shown in red font. **A**. ANXA2-miR-603. **B**. ANXA2-mutant Y23 (A23)-miR-603.

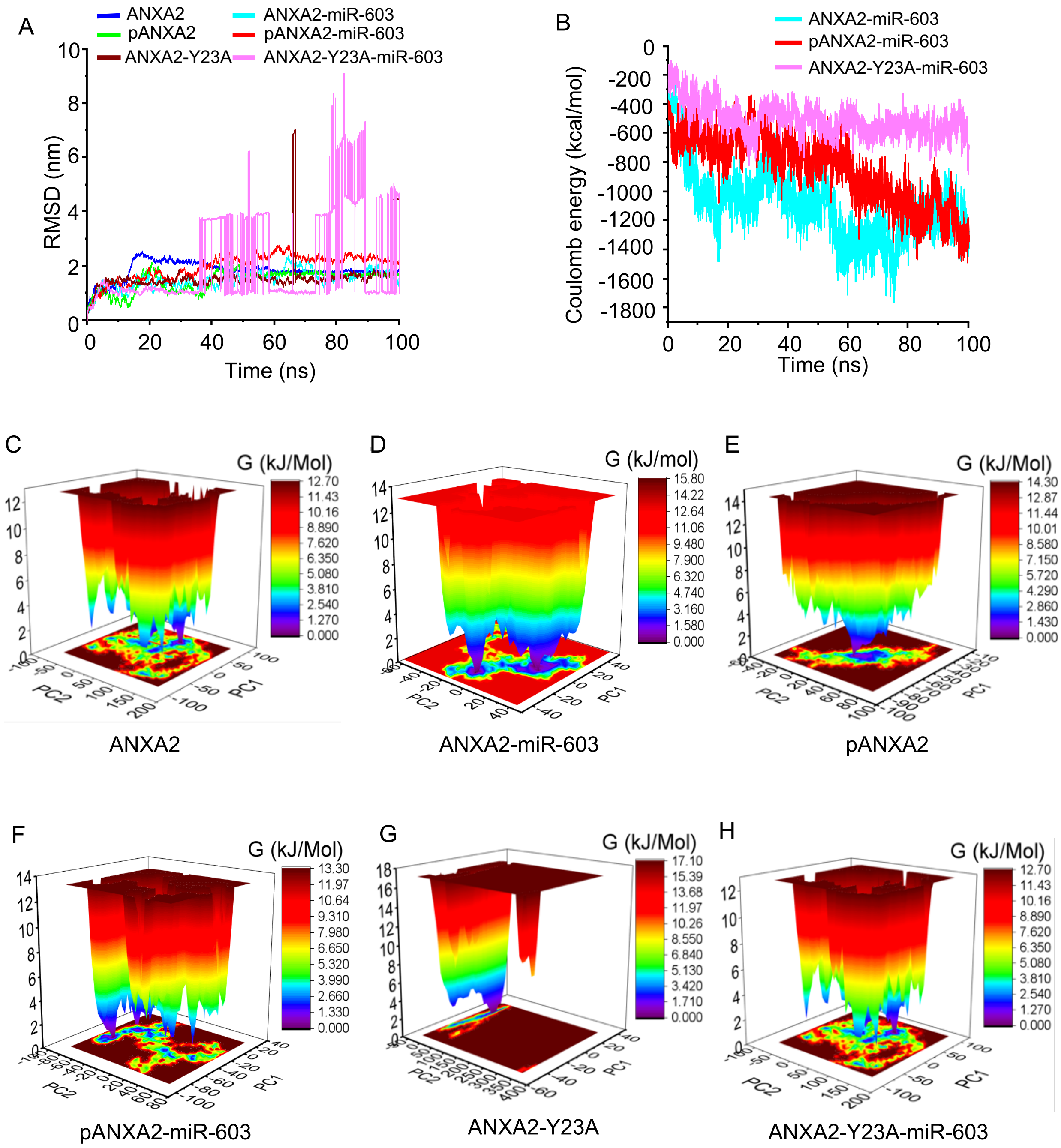


**Figure S10. In silico analysis of ANXA2-miR-603 interaction and complex stability.**

Molecular dynamics (MD) simulation-based energetic and stability analysis of ANXA2-miR-603 complexes. **A.** Root mean square deviation (RMSD) analysis of ANXA2, pANXA2, and ANXA2-Y23A, and their respective complexes with miR-603. Color identification: ANXA2 (blue), pANXA2 (green), ANXA2-Y23A (brown), ANXA2–miR-603 (teal), pANXA2–miR-603 (red), and ANXA2-Y23A-miR-603 (pink). **B.** Coulombic interaction energy profiles for complexes formed between miR-603 and ANXA2, or pANXA2, or ANXA2-Y23A mutant. Color identification: ANXA2-miR-603 (teal), pANXA2-miR-603 (red), and ANXA2-Y23A-miR-603 (pink). **C–H**. Free energy landscape (FEL) projections derived from principal component analysis, depicting the conformational space sampled by ANXA2 and miR-603 complex: ANXA2 (**C**), ANXA2-miR-603 (**D**), pANXA2 (**E**), pANXA2-miR-603 (**F**), ANXA2-Y23A (**G**), ANXA2-Y23A-miR-603 (**H**).

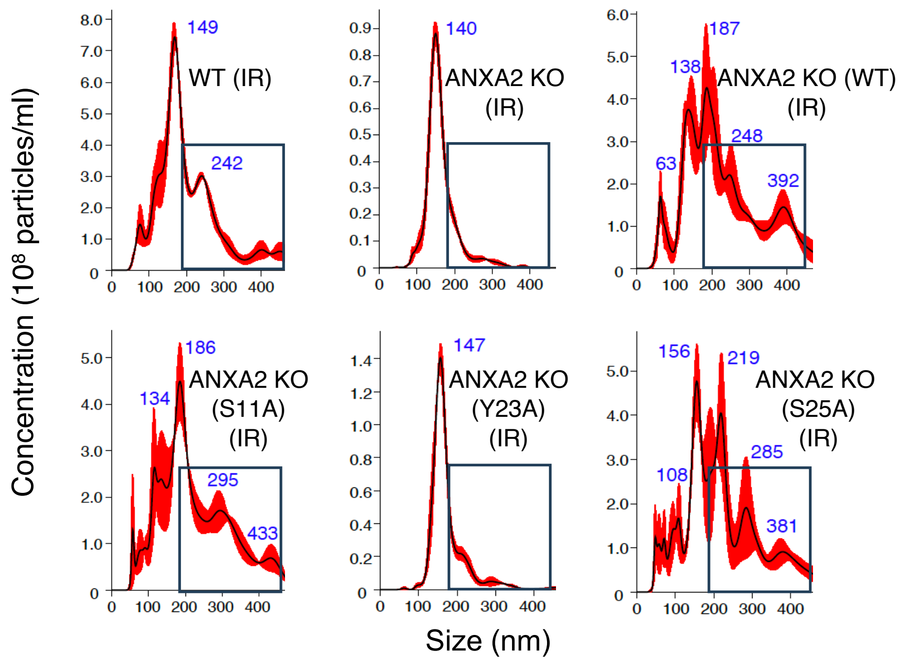


**Figure S11. Nanoparticle tracking analysis of MVs secreted from ANXA2 mutants expressing cells.**

Nanoparticle tracking analysis (NTA) of MVs isolated by differential centrifugation from ANXA2 mutants expressing conditioned media. Representative NTA traces illustrate particle size distribution, and quantification shows total particle concentration (particles/mL) and modal particle size (nm) of indicated samples.

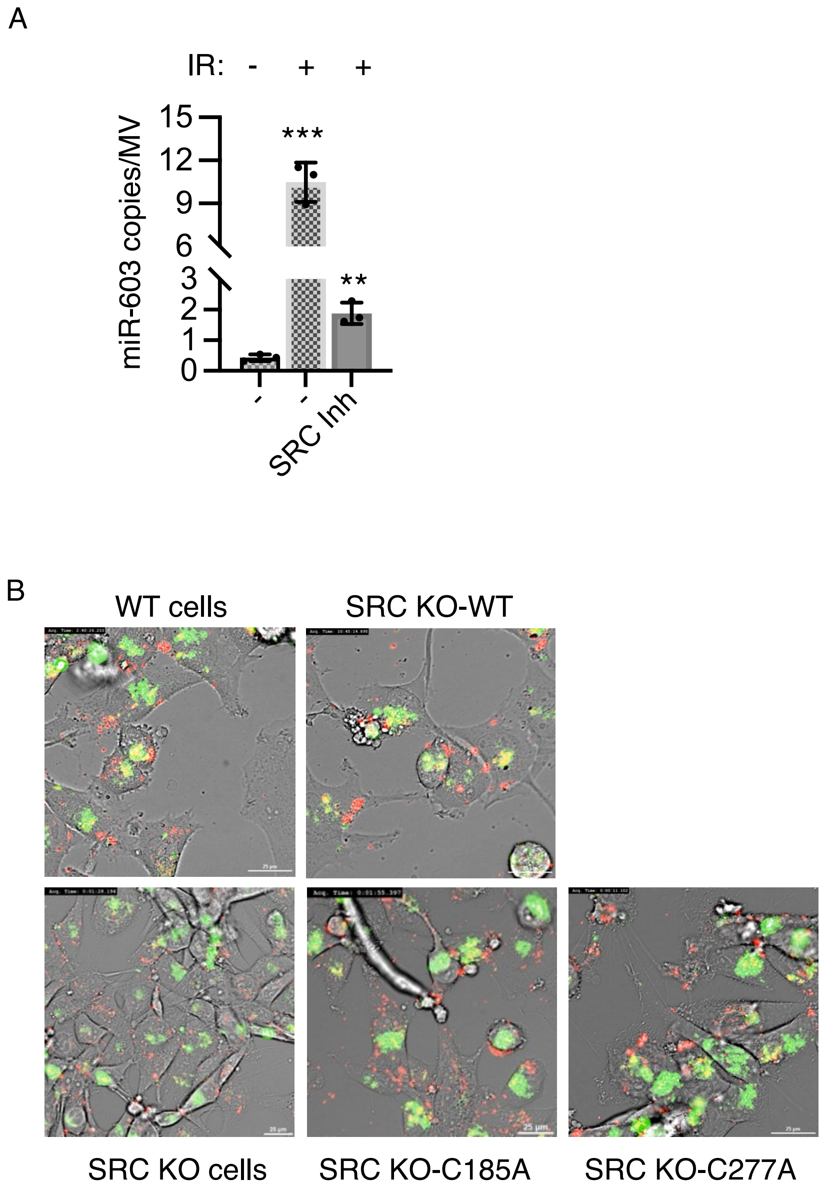


**Figure S12. SRC inhibition abolishes miR-603 export.**

**A.** Inhibition of SRC activity reduces packaging of miR-603 into microvesicles (MVs). LN340 cells were treated with the SRC inhibitor A-419259 trihydrochloride for 24 h, followed by exposure to ionizing radiation (IR). MVs were isolated from conditioned media, RNA was extracted, and miR-603 levels were quantified by RT-qPCR. **B.** LN340 WT and SRC KO cells stably expressing CD63-GFP transfected with Cy5-miR-603. The cells were washed, treated with 6 Gy ionizing radiation (IR) subjected to live-cell imaging. Representative live-cell videos are presented. *Top left*: WT (6 Gy); *bottom left*: SRC KO (6 Gy). *Top right*: SRC KO cells transfected with SRC WT plasmid (6 Gy); *bottom middle*: SRC KO cells transfected with SRC C185A plasmid (6 Gy); *bottom right*: SRC KO cells transfected with SRC C277A plasmid (6 Gy). Scale bar is 25 µm.

**Table S1. Oligonucleotide sequences of primers used in the study.**

| Name | Gene | Sequence |
|---|---|---|
| GS1 | ANXA2 FL sense | 5’ ATG TCT ACT GTT CAC GAA ATC CTG T 3’ |
| GS2 | ANXA2 FL antisense | 5’ GTC ATC TCC ACC ACA CAG GTA CAG C 3’ |
| GS3 | ANXA2 qP sense | 5’ CCAGGAGCTGCAGGAAATTA 3’ |
| GS4 | ANXA2 qP antisense | 5’ GTCACCAGATGTGTCCGAAATA 3’ |
| GS5 | ANXA2 (Mlu I) sense | 5’ AAA ACG CGT ATG TCT ACT GTT CAC GAA ATC CTG T 3’ |
| GS6 | ANXA2 (Xho I) antisense | 5’ AAAA CTC GAG GTC ATC TCC ACC ACA CAG GTA CAG C 3’ |
| GS7 | GAPDH sense | 5’ ACC CAG AAG ACT GTG GAT GG 3’ |
| GS8 | GAPDH antisense | 5’ TTC TAG ACG GCA GGT CAG GT 3’ |
| GS9 | ANXA2 S11A sense | 5’ TGC AAG CTC GCC TTG GAG GGT 3’ |
| GS10 | ANXA2 S11A antisense | 5’ ACC CTC CAA GGC GAG CTT GCA 3’ |
| GS11 | ANXA2 Y23A sense | 5’ CCA AGT GCA GCT GGG TCT GTC 3’ |
| GS12 | ANXA2 Y23A antisense | 5’ GAC AGA CCC AGC TGC ACT TGG 3’ |

| | | |
|---|---|---|
| GS13 | ANXA2 S25A sense | 5’ GCA TAT GGG GCT GTC AAA GCC 3’ |
| GS14 | ANXA2 S25A antisense | 5’ GGC TTT GAC AGC CCC ATA TGC 3’ |
| GS15 | SRC qP sense | 5’ CAG GCT GAG GAG TGG TAT TT 3’ |
| GS16 | SRC qP antisense | 5’ CCT TTC GTG GTC TCA CTT TCT 3’ |
| GS17 | SRC C185A sense | 5’ CGA AAG GTG CCTACG CCCTCT CAG TGT CTG AC 3’ |
| GS18 | SRC C185A antisense | 5’ GTC AGA CAC TGA GAG GGC GTA GGC ACC TTT CG 3’ |
| GS19 | SRC C277A sense | 5’ CTG GGC CAG GGC GCC TTT GGC GAG GTG 3’ |
| GS20 | SRC C277A antisense | 5’ CAC CTC GCC AAA GGC GCC CTG GCC CAG 3’ |
| GS21 | S100A10 qP sense | 5’ ACA AAG GAG GAC CTG AGA GTA 3’ |
| GS22 | S100A10 qP antisense | 5’ GGT CCG GGT CCT TCA TTA TTT 3’ |

**Methods**

**Molecular docking**

ANXA2 and miR-603 docking was conducted using the HDOCK web server (http://hdock.phys.hust.edu.cn/), which integrates template-based modelling and free docking approaches. Prior to docking, all crystallographic water molecules and heteroatoms were removed using Maestro. Docking study were performed between miR-603 and ANXA2 (WT), pANXA2, and the ANXA2-Y23A mutant. Residue Tyr23 in chain A was specified as the key binding-site residue to guide docking. For each system, the highest-ranked docking pose based on docking score and confidence score was selected for subsequent molecular dynamics simulations.

**Molecular dynamics simulations**

All-atom molecular dynamics (MD) simulations were performed using GROMACS version 2023.3. Simulations were conducted for unliganded ANXA2 (WT), pANXA2, and Y23A mutant proteins, as well as their corresponding protein-RNA complexes (ANXA2-miR-603, pANXA2-miR-603, and ANXA2-Y23A-miR-603). Prior to simulation, all non-essential heteroatoms, ligands, ions, and crystallographic water molecules were removed. Missing hydrogen atoms were added using Avogadro, and protonation states were assigned to reflect physiological pH using GROMACS utilities. Topology files for proteins and protein-miRNA complexes were generated using the AMBER99SB-ILDN force field for proteins and AMBER94 parameters for RNA to ensure force-field compatibility. Each system was solvated in a dodecahedral simulation box using the TIP3P water model, maintaining a minimum distance of 1.0 nm between the solute and box edges. Charge neutrality and physiological ionic strength (0.15 M) were achieved by replacing water molecules with $Ca^{2+}$ and/or $Cl^{-}$ ions using the gmx genion tool. Energy minimization was followed by equilibration under NVT conditions for 5 nano seconds (ns) and NPT conditions for an additional 5 ns. Temperature was maintained at 300 K using the V-rescale thermostat, and pressure was controlled at 1 bar using the Parrinello-Rahman barostat. Production MD simulations were carried out for 100 ns with a 2 femtosecond (fs) time step under periodic boundary conditions. Long-range electrostatic interactions were treated using the Particle Mesh Ewald (PME) method, and all hydrogen-containing bonds were constrained using the LINCS algorithm.

**Trajectory analysis and binding free energy calculations**

Post-simulation analysis were performed using standard GROMACS tools. Structural stability and flexibility were assessed by calculating root mean square deviation (RMSD), root mean square fluctuation (RMSF), and radius of gyration (Rg). Solvent-accessible surface area (SASA), hydrogen bonding patterns, and non-bonded interaction energies (Lennard-Jones and Coulombic energies) were quantified to evaluate protein-RNA interaction stability. Free energy landscapes

(FELs) were constructed to examine dominant conformational states sampled during the simulations. Binding free energies and per-residue energy contributions were estimated using the MM-PBSA approach. Structural visualization and trajectory inspection were performed using PyMOL and Visual Molecular Dynamics (VMD). Graphical analysis and plots were generated using Origin software.
